# Small molecule-mediated insulin hypersecretion induces transient unfolded protein response and loss of beta cell function

**DOI:** 10.1101/2022.03.07.482881

**Authors:** Karina Rodrigues-dos-Santos, Gitanjali Roy, Derk D. Binns, Magdalena G. Grzemska, Luiz F. Barella, Fiona Armoo, Melissa K. McCoy, Andrew V. Huynh, Jonathan Z. Yang, Bruce A. Posner, Melanie H. Cobb, Michael A. Kalwat

## Abstract

Pancreatic islet beta cells require a fine-tuned ER stress response for normal function; abnormal ER stress contributes to diabetes pathogenesis. Here, we identified a small molecule, SW016789, with time-dependent effects on beta cell ER stress and function. Acute treatment with SW016789 potentiated nutrient-induced calcium influx and insulin secretion, while chronic exposure to SW016789 transiently induced ER stress and shut down secretory function in a reversible manner. Distinct from the effects of thapsigargin, SW016789 did not affect beta cell viability or apoptosis, potentially due to a rapid induction of adaptive genes, weak signaling through the eIF2α kinase PERK, and lack of oxidative stress gene *Txnip* induction. We determined that SW016789 acted upstream of voltage-dependent calcium channels (VDCCs) and potentiated nutrient- but not KCl-stimulated calcium influx. Measurements of metabolomics, oxygen consumption rate, and G protein-coupled receptor signaling did not explain the potentiating effects of SW016789. In chemical co-treatment experiments we discovered synergy between SW016789 and activators of protein kinase C (PKC) and VDCCs, suggesting involvement of these pathways in the mechanism of action. Finally, chronically elevated calcium influx was required for the inhibitory impact of SW016789, as blockade of VDCCs protected human islets and MIN6 beta cells from hypersecretion-induced dysfunction. We conclude that beta cells undergoing this type of pharmacological hypersecretion have the capacity to suppress their function to mitigate ER stress and avoid apoptosis. These results have the potential to uncover beta cell ER stress mitigation factors and add support to beta cell rest strategies to preserve function.

## INTRODUCTION

Pancreatic islet beta cells sense and control circulating glucose concentrations by secreting insulin. In type 1 diabetes (T1D), autoimmune attack on beta cells results in their death and thus a lack of insulin. In type 2 diabetes (T2D), beta cells initially compensate for insulin resistance but eventually fail to secrete sufficient insulin to maintain euglycemia [1]. Interventions are aimed at restoring beta cell functionality, lowering insulin resistance, and reducing hyperglycemia. However, existing therapeutics have limitations, necessitating identification of new targets and pathways.

Beta cells sense elevated circulating blood glucose concentrations via the glucose transporters GLUT1/3 [2]. Upon cellular entry, glucose metabolism via glycolysis and the tricarboxylic acid cycle yields a relative increase in [ATP/ADP], causing ATP-sensitive potassium (K_ATP_) channel closure. This closure reduces the plasma membrane permeability to potassium, causing membrane depolarization, opening of voltage-dependent Ca^2+^ channels (VDCCs) and Ca^2+^ influx [3]. Ca^2+^ influx also stimulates oxidative phosphorylation in the mitochondria, driving a metabolic cycle that sustains ATP production and insulin secretion [4, 5]. Ca^2+^ influx triggers the fusion of insulin secretory vesicles with the plasma membrane resulting in insulin exocytosis [6]. Nutrient metabolism further enhances secretion without further Ca^2+^ influx via the amplifying pathway [7–9]. To handle their specialized secretory function, beta cells engage the unfolded protein response of the endoplasmic reticulum (UPR^ER^) [10]. The UPR^ER^ is crucial in beta cells, as its disruption (e.g., PERK loss-of-function) leads to beta cell failure and diabetes [11]. Conversely, unmitigated UPR^ER^ is also detrimental to beta cell function and survival, eventually leading to induction of pro-apoptotic transcription factors including *Ddit3* (CHOP) and *Txnip* [12, 13].

Many studies have elucidated aspects of beta cell biology that may be targeted to improve overall beta cell health and function, including the UPR^ER^, beta cell dedifferentiation, and the role of immature beta cells in whole islet function [14–18]. However, the high-throughput identification of novel small molecules that alter pancreatic beta cell function has lagged in comparison. Here, we used a secreted insulin-luciferase-based high-throughput screening tool to identify a small molecule, SW016789, which acutely and chronically alters beta cell secretory responses [19, 20]. We hypothesized the chronic effects on function were a result of acute hypersecretion which induced stress pathways (e.g., ER stress) that led to a shutdown of beta cell function, but not beta cell death. We investigated the mechanisms of action of SW016789 using a multipronged approach including measuring Ca^2+^ influx and insulin secretion; metabolomics and mitochondrial function; gene expression; and co-treatment with compounds that may interfere with the effects of SW016789. Our results with SW016789 agree with a model of beta cell plasticity wherein the effects of persistent secretory stress are mitigated to avoid cell death at the expense of function. Chemical tools like SW016789 present an opportunity to discover previously unrecognized or poorly characterized genes and pathways involved in these processes in the beta cell.

## METHODS

### Antibodies and Reagents

Source information for all small molecules and antibodies used in this work are listed in electronic supplementary material (ESM) **Table S4**.

### Small molecule screening

High-throughput screening of small molecules in Ins-GLuc MIN6 cells was as described [21]. Briefly, cells were seeded at ∼15e6 cells per T175 flask and grown 7 days until confluence, then trypsinized, resuspended in medium, passed through a 40 µm cell strainer and diluted to 1.5e6 cells/ml. Cells were dispensed at 50 µl per well in 384-well opaque white cell culture dishes using a BioTek Multiflo FX liquid handler to yield 7.5e4 cells/well. Plates were placed in a tissue culture incubator for 72 h. Next, 0.5 µl each of DMSO (negative control, 1%), thapsigargin (positive control, 1 mM), and small molecules (5 µM) were added robotically to the plates and the cells were incubated 24 h. On the day of the assay, using a BioTek Multiflo FX liquid handler, cells were washed twice with KRBH (75 µL then 50 µL). To remove medium and buffer between washes, plates were centrifuged upside-down in collection trays at 30 x g for 1 min. Cells were then incubated in 25 µl of KRBH containing 200 mM diazoxide for 60 min. Next, 25 µl of 2X stimulation buffer in KBRH was added to yield a final concentration of 20 mM glucose, 35 mM KCl and 250 mM diazoxide in a total volume of 50 µl. The plates were incubated for 60 min at 37°C in a tissue culture incubator. Afterward, 20 µl of GLuc assay working solution (5 mM KCl, 15 mM Hepes pH 7.4, 24 mM NaHCO3, 1 mM MgCl2, 2 mM CaCl2, 300 mM sodium ascorbate, and 3.54 µM coelenterazine) was added to each well. Within 10-20 min, the luminescence from each well of the plates was measured using a Perkin Elmer EnVision multi-mode plate reader with 0.1 sec integration time. The raw data were corrected for plate effects using a GeneData proprietary algorithm and then each value on a per plate basis was subjected to either single-point or two-point normalization, followed by calculation of the robust Z (RZ) score for each well. Hit compounds with an |RZ|>3 were selected for structural clustering analysis. Representative compounds were then cherry picked and confirmed in triplicate in the same InsGLuc-MIN6 secretion assay and tested for toxicity in MIN6 cells by Cell Titer Glo assay after 72 h exposure.

### Cell culture and treatments

Culture of MIN6 beta cells has been described [20]. MIN6 cells were cultured in high glucose DMEM containing 10% FBS, 50 µM beta-mercaptoethanol, 1 mM pyruvate, 100 U/ml penicillin, and 100 μg/ml streptomycin. Prior to glucose stimulation experiments (as in **Fig. 3a-d**), cells were washed twice with and incubated for 2 h in freshly prepared glucose-free modified Krebs-Ringer bicarbonate (KRBH) buffer (5 mM KCl, 120 mM NaCl, 15 mM HEPES, pH 7.4, 24 mM NaHCO_3_, 1 mM MgCl_2_, 2 mM CaCl_2_, and 1 mg/ml radioimmunoassay-grade BSA). MIN6 cells were stimulated with glucose as indicated and secreted insulin and insulin content were measured using HTRF assays (Cisbio). To generate lysates for immunoblotting, cells were lysed in 1% NP40 lysis buffer (25 mM HEPES, pH 7.4, 1% Nonidet P-40, 10% glycerol, 137 mM NaCl, 1 mM EDTA, 1 mM EGTA, 50 mM sodium fluoride, 10 mM sodium pyrophosphate, 1 mM sodium orthovanadate, 1 mM phenylmethylsulfonyl fluoride, 10 μg/ml aprotinin, 1 μg/ml pepstatin, 5 μg/ml leupeptin), rotated (10 min, 4°C), centrifuged (10,000 x g; 10 min; 4°C), and stored at -80°C. InsGLuc-MIN6 cells were generated and characterized as previously described [20] and were determined to be negative for mycoplasma. Mouse intestinal L cell line GLUTag was originally generated by Daniel Drucker (Mount Sinai Hospital) and were obtained from the lab of George G. Holz (Upstate Medical University). GLUTag cells were cultured in low glucose DMEM with 10% FBS and 100 U/ml penicillin, 100 μg/ml streptomycin. The human somatostatinoma line QGP-1 was provided by Dawn E. Quelle (University of Iowa). QGP-1 cells were cultured in RPMI-1640 with 10% FBS, 1 mM pyruvate, 100 U/ml penicillin, and 100 μg/ml streptomycin. GLUTag and QGP-1 cells were plated in 12 well dishes and treated in medium as indicated in the figure legend before harvesting for RNA purification and downstream gene expression analysis.

### Secreted Gaussia luciferase assays

InsGLuc-MIN6 cells were plated in 96-well dishes at 1e5 cells/well and cultured for 3-4 days before compound treatment either overnight (24 h) in medium or acutely (1 h) in KRBH. Treatments in 96-well assays were performed with at least three replicate wells and in at least three independent passages of cells. For assays, cells were washed twice with KRBH and preincubated in 100 µl of KRBH (250 µM diazoxide where indicated) for 1 h. The solution was then removed, and cells were washed once with 100 µl of KRBH and then incubated in KRBH with or without the indicated stimulation and compound treatments for 1 h. 50 µl of KRBH was collected from each well and pipetted into a white opaque 96-well plate Optiplate-96 (Perkin-Elmer). Fresh GLuc assay working solution was then prepared by adding coelenterazine stock solution into assay buffer to a final concentration of 10 µM. 50 µl of working solution was then rapidly added to the wells using an Integra Voyager 1250 µl electric multi-channel pipette for a final concentration of 5 µM coelenterazine. After adding reagent to the plate, plates were spun briefly, and luminescence was measured on a Synergy H1M2 plate reader (BioTek). The Gen5 software protocol was set to shake the plate orbitally for 3 seconds and then read the luminescence of each well (integration, 100 ms; gain,150).

### Human islet culture, drug treatments and static culture secretion assays

Cadaveric human islets were obtained through the Integrated Islet Distribution Program (IIDP). Islets were isolated by the affiliated islet isolation center and cultured in PIM medium (PIM-R001GMP, Prodo Labs) supplemented with glutamine/glutathione (PIM-G001GMP, Prodo Labs), 5% Human AB serum (100512, Gemini Bio Products), and ciprofloxacin (61-277RG, Cellgro, Inc) at 37°C and 5% CO2 until shipping at 4°C overnight. Human islets were cultured upon receipt in complete CMRL-1066 (10% FBS, 100 U/ml penicillin, 100 μg/ml streptomycin). For drug treatments and static culture, human islets (50-75) were hand-picked under a dissection microscope and transferred to low-binding 1.5 ml tubes (50 islets/tube). Islets were cultured 24 h in 500 µL of complete CMRL-1066 medium containing indicated drug treatments. For static culture insulin secretion experiments, the islets were washed twice with 500 µL of KRBH (134 mM NaCl, 4.8 mM KCl, 1 mM CaCl_2_, 1.2 mM MgSO_4_, 1.2 mM KH_2_PO_4_, 5 mM NaHCO_3_, 10 mM HEPES pH 7.4, 0.1% radioimmunoassay-grade BSA) and preincubated in KRBH (500 µL, 1 mM glucose; 1 h) before switching to 500µL KRBH containing either 1 or 16 mM glucose for 1 h. Supernatants were collected, centrifuged (10,000 x g; 5 min) and transferred to fresh tubes for storage at -80°C. Total insulin content was extracted by acid-ethanol (1.5% HCl in 70% ethanol) overnight at -80°C and the solution was neutralized with an equal volume of 1 M Tris pH 7.4 prior to insulin assays. Secreted insulin and insulin content were measured using HTRF assays. Human islet characteristics and donor information are listed in **ESM Human Islet Checklist**.

### Cell death and viability assays

To measure live and dead cells as well as the live/dead cell ratio, MultiTox-Fluor (Promega) was used according to the manufacturer’s instructions. Briefly, MIN6 cells were seeded into 96 well black-walled clear bottom tissue culture dishes at 5e4-1e5 cells/well and culture for 3-5 d before treatment with compounds as indicated in the figure legend. On the day of the assay, cell-permeant GF-AFC substrate (live cell readout) and cell-impermeant bis-AAF-R110 substrate (dead cell readout) reagents were diluted into kit assay buffer as a 2X working solution. 100 µL of 2X working solution was added to each well of cells in 100 µL of medium and incubated 30 min at 37°C. Plates were read on a BioTek Synergy H1M2 plate reader (400_Ex_/505_Em_ for live cells; 485_Ex_/520_Em_ for dead cells). Digitonin was a positive control for cell death which lysed cells immediately prior to addition of 2X working solution. Viability and death were normalized to DMSO control.

### Ca^2+^ influx assays

Ca^2+^ influx measurements were completed in MIN6 cells plated in triplicate in black-walled clear-bottom 96-well tissue culture imaging plates. Cells were preincubated for 1 h in glucose-free KRBH, followed by loading with Fura-2-LR-AM (5 µM) for 25 min and a 5 min washout. Using a BioTek Synergy H1 plate reader equipped with dual monochromators, relative Ca^2+^ levels were measured (Ex_340/380_/Em_510_ gain=150). Recordings included 5 min baseline followed by 5-30 min post-addition of stimulation as indicated. Data were normalized by subtracting the mean of the respective baseline reads. Area under the curves (AUCs) were calculated from peaks detected in GraphPad Prism using a baseline of 0 ΔRFU, restricted to peaks that go above baseline by more than 0.05 ΔRFU and defined by at least 10 adjacent points.

### Immunoblotting

40-50 µg of cleared cell lysates were separated on 10% gels by SDS-PAGE and transferred to nitrocellulose for immunoblotting. All membranes were blocked in Odyssey blocking buffer (Licor) for 1 h before overnight incubation with primary antibodies diluted in blocking buffer (**ESM Table 4**). After three 10 min washes in TBS-T (20 mM Tris-HCl pH 7.6, 150 mM NaCl, 0.1% Tween-20), membranes were incubated with fluorescent secondary antibodies for 1 h at room temperature. After three 10 min washes in TBS-T, membranes were imaged on a Licor Odyssey scanner.

### Relative gene expression

RNA was isolated from MIN6 cells using the PureLink RNA Mini Kit (Life Technologies) or the Aurum Total RNA Mini kit (Bio-Rad). Briefly, after indicated treatments, medium was removed from cells and lysis buffer (with β-mercaptoethanol) was added to cells. Cells were scraped, lysates transferred to 1.5 mL tubes on ice and then transferred to -80°C for storage. Samples were processed according to the kit manufacturer’s instructions, including on-column digestion with RNase-free DNase. RNA concentration was measured using an Implen Nanophotometer or Nanodrop spectrophotometer and verified to have A_260/280_ ratios >2.0. 1000 ng of RNA was converted into cDNA using the iScript cDNA synthesis kit (Bio-Rad) following manufacturer instructions and the resulting cDNA was diluted 20-fold with water. One μl of diluted cDNA was used in 10 μl qPCR reactions using 2X SYBR Bio-Rad master mix and 250 nM of each primer. Reactions were run in 384-well format on a CFX384 (Bio-Rad) or QuantStudio 5 (Thermo). qPCR data was analyzed using CFX Manager (Bio-Rad) with 18S RNA as the reference gene. Relative expression was calculated by the 2^-ΔΔCt^ method. All primer details are listed in ESM **Table 4.**

### Bioenergetics

Oxygen consumption rates were measured using a Seahorse XFe96 (Agilent). MIN6 cells were seeded at a density of 5e4 cells/well in XF96 cell culture microplates and cultured for 48 h. On the day before the assay a Seahorse cartridge was hydrated with calibrant solution and incubated overnight. On the day of the assay, the XFe96 PrepStation was set to 37°C and hydrated cartridge was placed inside. MIN6 cells were washed twice with Seahorse-modified Krebs-Ringer HEPES buffer (S-KRH: 5 mM KCl, 114 mM NaCl, 20 mM Hepes pH 7.4, 1.2 mM MgSO_4_, 2.5 mM CaCl_2_, 1.2 mM KH_2_PO_4_ and pH carefully adjusted to 7.4, BSA and NaHCO_3_ are not present) and placed in the PrepStation to equilibrate for 1 h. The hydrated Seahorse cartridge was placed on an empty utility plate to be loaded with compounds diluted in S-KRH. The drug cartridge was returned to the utility plate with calibrant solution and inserted into the XFe96 for calibration. After calibration, the cell culture plate was inserted into the XFe96 and MIN6 cells were monitored for three 3 min cycles to measure baseline oxygen consumption (OCR) and extracellular acidification rates (ECAR). Stimulations were injected as indicated and after each stimulation cells were monitored for three 3 min cycles. Data were analyzed in the Seahorse WAVE software (version 2.61.53).

### Metabolomics

Metabolomic analysis was performed as described [22, 23]. MIN6 cells grown to confluence in 6 well dishes were preincubated in glucose-free KRBH for 2 h at 37°C. DMSO (0.1%), SW016789 (5 µM), and/or glucose (20 mM) were added as indicated and cells were incubated a further 30 min at 37°C. After indicated treatments, cells were washed with 1 ml ice cold saline and harvested in 1 ml of 80% methanol at 80°C. Cells were scraped and transferred to 1.5 ml tubes for three freeze-thaw cycles in liquid nitrogen. Lysates were clarified at 14,000 rpm for 15 min at 4°C. Metabolite containing supernatants were transferred to a new 1.5 ml tube and lyophilized using a SpeedVac. Breathable membranes were placed on top of the open tubes to prevent cross contamination and the SpeedVac was operated without heat for 4 h. Pellets were stored at 80°C prior to analysis by the Metabolomics Facility at Children’s Medical Center Research Institute at UT Southwestern (Dallas, TX). Metabolites were reconstituted in 100 ml of 0.03% formic acid in analytical grade water, vortex mixed, and centrifuged to remove debris. Thereafter, the supernatant was transferred to a high-performance liquid chromatography (HPLC) vial for the metabolomics study. Targeted metabolite profiling was performed using a liquid chromatography mass spectrometry/mass spectrometry (LC/MS/MS) approach. Separation was achieved on a Phenomenex Synergi Polar RP HPLC column (150 x 2 mm, 4 µm, 80 Å) using a Nexera Ultra High Performance Liquid Chromatograph (UHPLC) system (Shimadzu Corporation). The mobile phases employed were 0.03% formic acid in water (A) and 0.03% formic acid in acetonitrile (B). The gradient program was as follows: 0-3 min, 100% A; 3-15 min, 100% - 0% A; 15-21 min, 0% A; 21-21.1 min, 0% - 100% A; 21.1-30 min, 100% A. The column was maintained at 35°C and the samples kept in the autosampler at 4°C. The flow rate was 0.5 ml/min, and injection volume 10 µl. The mass spectrometer was an AB QTRAP 5500 (Applied Biosystems SCIEX) with electrospray ionization (ESI) source in multiple reaction monitoring (MRM) mode. Sample analysis was performed in positive/negative switching mode. Declustering potential (DP) and collision energy (CE) were optimized for each metabolite by direct infusion of reference standards using a syringe pump prior to sample analysis. The MRM MS/MS detector conditions were set as follows: curtain gas 30 psi; ion spray voltages 5000 V (positive) and 1500 V (negative); temperature 650°C; ion source gas 150 psi; ion source gas 250 psi; interface heater on; entrance potential 10 V. Dwell time for each transition was set at 3 msec. Samples were analyzed in a randomized order, and MRM data was acquired using Analyst 1.6.1 software (Applied Biosystems SCIEX). Chromatogram review and peak area integration were performed using MultiQuant software version 2.1 (Applied Biosystems SCIEX). Although the numbers of cells were similar and each sample was processed identically and randomly, the peak area for each detected metabolite was normalized against the protein content of that sample to correct any variations introduced from sample handling through instrument analysis. The normalized area values were used as variables for the multivariate and univariate statistical data analysis. The chromatographically co eluted metabolites with shared MRM transitions were shown in a grouped format, i.e., leucine/isoleucine. Principle component analysis (PCA) was completed using SIMCA-P software (version 13.0.1, Umetrics). To generate heat maps, metabolite TIC values were normalized to the average of ‘DMSO Basal’ and expressed in log_2_. Heatmap was generated using MetaboAnalyst 5.0 [24] using Euclidean distance with the Ward clustering algorithm, without autoscaling of the features. Statistical differences were assessed by one-way ANOVA with Fisher’s LSD post-hoc test with an FDR <0.05 in MetaboAnalyst 5.0. Raw TIC values and table of significantly altered metabolites are provided in **ESM Table 3**.

### Statistical Analysis

Quantitated data are expressed as mean ± SD. Data were evaluated using one- or two-way ANOVA as indicated and considered significant if P < 0.05. Graphs were made in GraphPad Prism 9 and figures assembled in Affinity Designer (Serif).

## RESULTS

### High-throughput screen identifies small molecule modulator of insulin secretion

In order to discover small molecules that alter beta cell function, we utilized the insulin-Gaussia luciferase (GLuc) fusion reporter expressed in MIN6 beta cells (InsGLuc-MIN6) [20]. InsGLuc is processed and leads to co-secretion of Gaussia luciferase (GLuc) with insulin. Secreted GLuc activity in this manner has been well-established as a proxy for insulin release [19, 25–29]. Our approach involved treating InsGLuc-MIN6 beta cells for 24 h with compounds followed by a brief wash-out prior to testing their sustained impact on insulin secretion. This strategy probes a distinct phenotypic space and is complementary to results from studies utilizing acute treatments during stimulation [19]. We screened an 8,320 compound diversity subset approximately representing the chemical space of the >300,000 compound library at the UT Southwestern Medical Center High-Throughput Screening Core. InsGLuc-MIN6 beta cells were treated with control and test compounds for 24 h in normal medium followed by a washout and stimulation with glucose and KCl in the presence of diazoxide (**Fig. 1a**). Among the hits, chronic exposure to SW016789 caused inhibition of secretion in InsGLuc-MIN6 cells with an IC_50_ <1 µM (**Fig. 1b**) and was chosen for further study. Glucose-stimulated InsGLuc secretion was blunted in MIN6 cells chronically exposed to resupplied stocks of SW016789, validating screening results (**Fig. 1c**). Chronic pretreatment (24 h) of isolated human islets with SW016789 also impaired glucose-stimulated insulin secretion (GSIS) (**Fig. 1d,e**), without a significant impact on insulin content (**Fig. 1f**). SW016789 did not impact cell viability (**Fig. 1g**), cell death (**Fig. 1h**), or the live/dead cell ratio (**Fig. 1i**) in MIN6 cells treated for up to 72 h. During this time course, SW016789-treated cells remained unresponsive to stimulation with glucose or KCl (**Fig. 1j**), but were able to mount significant, although blunted, responses to glucose if combined with stimulators of cAMP generation or protein kinase C (PKC) activation (**Fig. 1j**). Beta cells exposed to SW016789 for 24 h recovered function after a 48 h washout (**Fig. 1k**), indicating the effects are reversible.

**Fig. 1.**
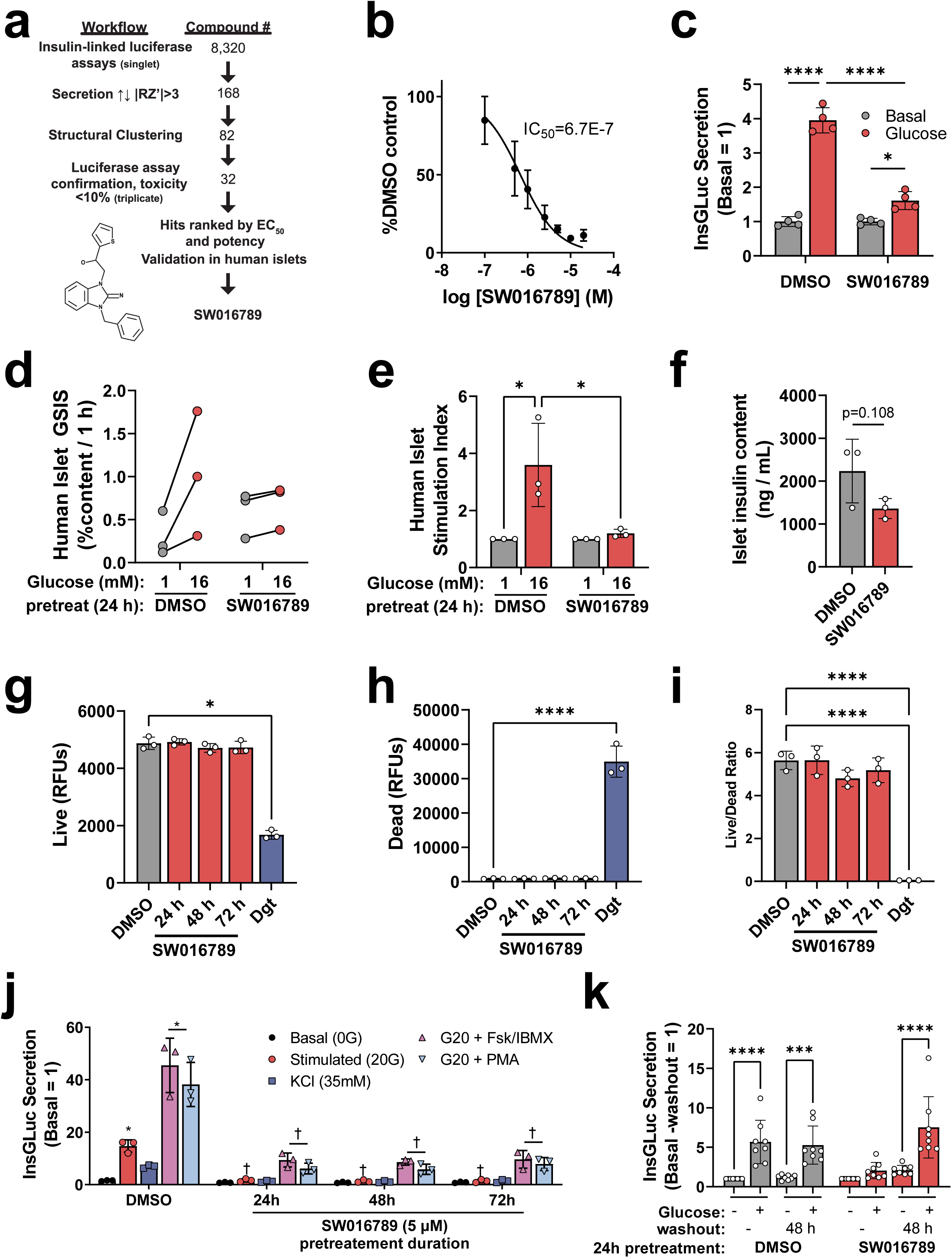
Discovery of SW016789 as a small molecule chronic suppressor of insulin secretion. **(a)** High-throughput screening workflow leading to the discovery of SW016789. Screening was performed on an 8,320-compound diversity subset representing the chemical space of the full compound library at the UT Southwestern Medical Center High-Throughput Screening Core. Cells were treated with control and test compounds for 24 h in normal medium followed by a washout and stimulation with glucose and KCl in the presence of diazoxide and finally subjected to InsGLuc assays. These maximal stimulatory conditions afforded the assay a large dynamic range (average Z-score >0.5) and resulted in 168 primary hits which after structural clustering were narrowed down to 82, of which 32 confirmed in triplicate. **(b)** Concentration response curve testing to determine EC_50_ of SW016789 InsGLuc-MIN6 after a 24 h exposure. Data are the mean ± SD of three independent assays. **(c,d)** In mouse InsGLuc-MIN6 beta cells (**c**) and human islets (**d**), 24 h exposure to SW016789 (5 µM) resulted in suppression of GSIS during assays in the absence of compound. Data are the mean ± SD of 3-4 independent experiments. **(g-i)** Exposure of MIN6 beta cells to DMSO (0.1%) or SW016789 (5 µM) for up to 72 h did not alter relative amounts of (**g**) live cells, (**h**) dead cells, or (**i**) the live/dead cell ratio. Digitonin (Dgt) treatment was a positive control for cell death added just prior to the assay. **(j)** InsGLuc-MIN6 cells were exposed to SW016789 (5 µM) for 24-72 h followed by a 1 h washout in glucose-free KRBH. Cells were then stimulated for 1 h with the indicated treatments (0G, glucose-free; 20G, glucose 20 mM; Fsk, forskolin 10 µM; IBMX 100 µM; PMA, phorbol myristate acetate 100 nM). *, P<0.05 vs respective Basal; †, P<0.05 vs respective DMSO. **(k)** InsGLuc-MIN6 cells were exposed to SW016789 (5 µM) for 24 h followed by a 48 h washout in complete medium. Cells were then assayed for glucose-stimulated InsGLuc secretion. All data are the mean ± SD of at least three independent experiments. *, P<0.05 by two-way ANOVA.

### SW016789 acutely potentiates nutrient-stimulated insulin secretion

We determined that exposure of InsGLuc-MIN6 cells to SW016789 for 4-8 h resulted in significant inhibition of GSIS while acute (≤2 h) exposure caused a marked potentiation of GSIS (EC_50_ = 2.7 µM) (**Fig. 2a, ESM Fig. 1a,b**). Structure-activity relationship studies revealed that while SW016789 is chiral, both enantiomers retain activity (**ESM Fig. 1c-f**) and the alcohol at the chiral center is important for activity (**ESM Fig. 1g**). SW016789 potentiated amino acid-induced secretion (**Fig. 2b**) but had no effect in cells treated with diazoxide and depolarizing KCl (**Fig. 2c**). However, SW016789 elicited a response from cells treated with diazoxide and glucose without elevated KCl. These data suggested the pathway activated by SW016789 requires nutrient metabolism and may impact Ca^2+^ influx. Accordingly, chronic SW016789 treatment suppressed glucose-stimulated Ca^2+^ influx (GSCa) (**Fig. 3a,b**), and acute SW016789 treatment potentiated nutrient- but not KCl-stimulated Ca^2+^ influx (**Fig. 3c-f**). Considering these results, we determined the effects of SW016789 on Ca^2+^ influx in the diazoxide paradigm. Beta cells were either pre-treated for 24 h or treated with compound simultaneously with stimulation. With chronic SW016789, stimulation with depolarizing KCl or KCl/glucose caused a rapid and sustained rise in Ca^2+^ influx which was significantly lower than control by ∼40% (**ESM Fig. 2a**), while acute SW016789 had no impact (**ESM Fig. 2b**). We also observed a slight increase in GSCa even in the presence of diazoxide and low KCl (**ESM Fig. 2c**), which was consistent with secretion data (**Fig. 2c**).

**Figure 2.**
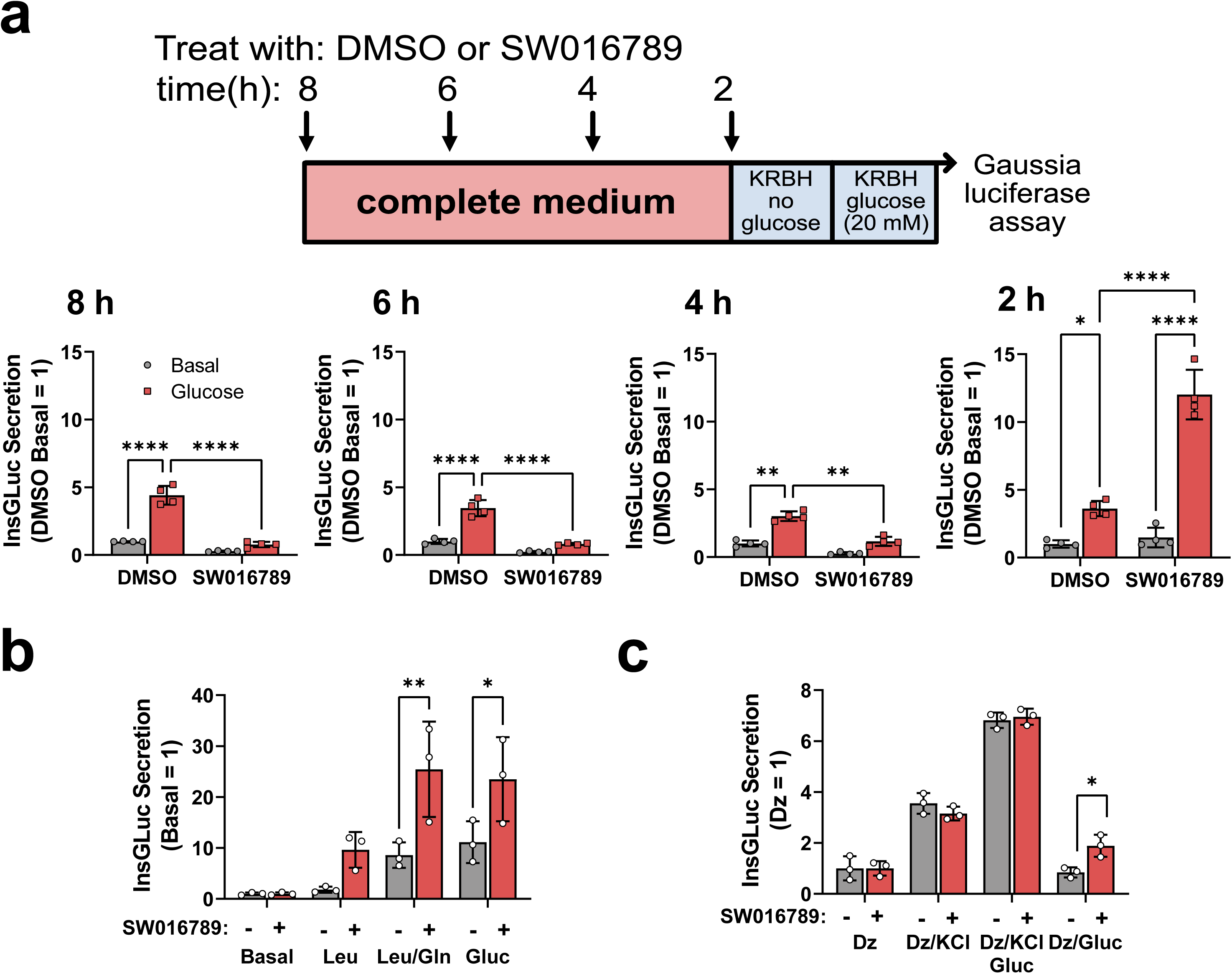
Time-course studies and phenotypic assays reveal SW016789 as an acute potentiator of nutrient-stimulated insulin secretion. (**a**) InsGLuc-MIN6 cells were treated with SW016789 (5 µM) in a time-course as indicated in the diagram. At the end of the time-course, glucose-stimulated InsGLuc secretion was assayed. Data are the mean ± SD of N=4. (**b**) InsGLuc-MIN6 cells were exposed to SW016789 (5 µM) for 1 h during stimulation with glucose (20 mM), leucine (5 mM), or leucine/glutamine (Leu/Gln; 5mM each) or (**c**) in the presence of diazoxide (250 µM) with or without KCl (35 mM) or glucose (20 mM). Data are the mean ± SD of N=3. *, P<0.05 by two-way ANOVA.

A potential source of the SW016789-enhanced Ca^2+^ influx is the endoplasmic reticulum (ER). We tested whether SW016789 impacted ER Ca^2+^ stores or store-operated Ca^2+^ entry (SOCE). MIN6 beta cells were pre-treated with SW016789 for 2, 6, and 24 h and subjected to Ca^2+^ assays including thapsigargin stimulation to block SERCA2, leading to an observable Ca^2+^ leak into the cytosol (**ESM Fig. 2d**). Treatment with SW016789 did not alter this indirect measure of ER Ca^2+^. Subsequently, Ca^2+^ was reintroduced to measure SOCE. Pre-exposure to SW016789 for 2-6 h caused a small, but significant, decrease in SOCE, but this effect was absent in cells treated for 24 h, suggesting that in beta cells exposed to SW016789, ER Ca^2+^ and SOCE responses were unmodified and may not underlie the acute potentiating or chronic inhibitory effects of SW016789. Additionally, acute SW016789 exposure did not impact muscarinic receptor-linked Gαq-stimulated Ca^2+^ influx (**ESM Fig. 2e**).

### SW016789 is unlikely to exert its beta cell effects via G protein-coupled receptors

Because of its acute potentiating activity, we considered SW016789 may act through a G protein-coupled receptors (GPCRs). Acute potentiation of insulin secretion can occur via activation of Gαs-linked GPCRs and resultant cAMP generation [30]. To monitor relative cAMP production, we used the BRET reporter CAMYEL stably expressed in MIN6 cells [31]. The positive control (forskolin+IBMX) caused a marked increase in cAMP, while acute stimulation with SW016789, with or without glucose, yielded no consistent alterations in signal (**ESM Fig. 3a,b**). To test a wider range of GPCRs we also submitted SW016789 for screening against the GPCR library at the NIMH Psychoactive Drug Screening Program at University of North Carolina [32] (**ESM Fig. 3c**). Overall, SW016789 exhibited little to no activity against the GPCRs in the panel, including well-known beta cell GPCRs (**ESM Table 1**). Only CXCR7 (ACKR3) out of 320 GPCRs was a substantial primary hit, however confirmation studies determined the EC_50_ of SW016789 on CXCR7 was ∼26 µM (**ESM Fig. 3d**), well above the EC_50_ for beta cell activity (**Fig. 1b, ESM Fig. 1a**). In line with this result, the CXCR7 agonist VUF11207 [33] had a negligible impact on InsGLuc secretion during chronic or acute treatments (**ESM Table 2**). These data suggested CXCR7 is unlikely to mediate the effects of SW016789 in beta cells where its activity is observed in the 1-5 µM range. Together, these data suggested SW016789 does not exhibit substantial off-target activity on GPCRs and that other mechanisms may be involved in its effects in beta cells.

### Beta cell metabolomic and mitochondrial responses are normal following acute SW016789 exposure

Because SW016789 required nutrient co-stimulation to elicit effects on Ca^2+^ influx, we hypothesized altered metabolism may contribute to the mechanism of action. To test this, we performed targeted metabolomics on normal MIN6 beta cells stimulated with or without glucose in the presence of DMSO or SW016789 for 30 min (**Fig. 4a**). The amplifying effect of SW016789 on GSIS was confirmed by insulin measurements and the cellular extracts were submitted for metabolomic analysis (**Fig. 4b, ESM Table 3**). Expectedly, glucose induced glycolysis and TCA intermediates and the reduced/oxidized glutathione ratio (**Fig. 4c,d**). In each of these cases, SW016789 had no impact. These results suggested SW016789 is unlikely to exert effects by acutely altering well-known metabolic pathways. Beta cell oxygen consumption rates and extracellular acidification rates were also unaffected by acute exposure to SW016789 as measured by Seahorse assays (**Fig. 4e-g**).

**Figure 3.**
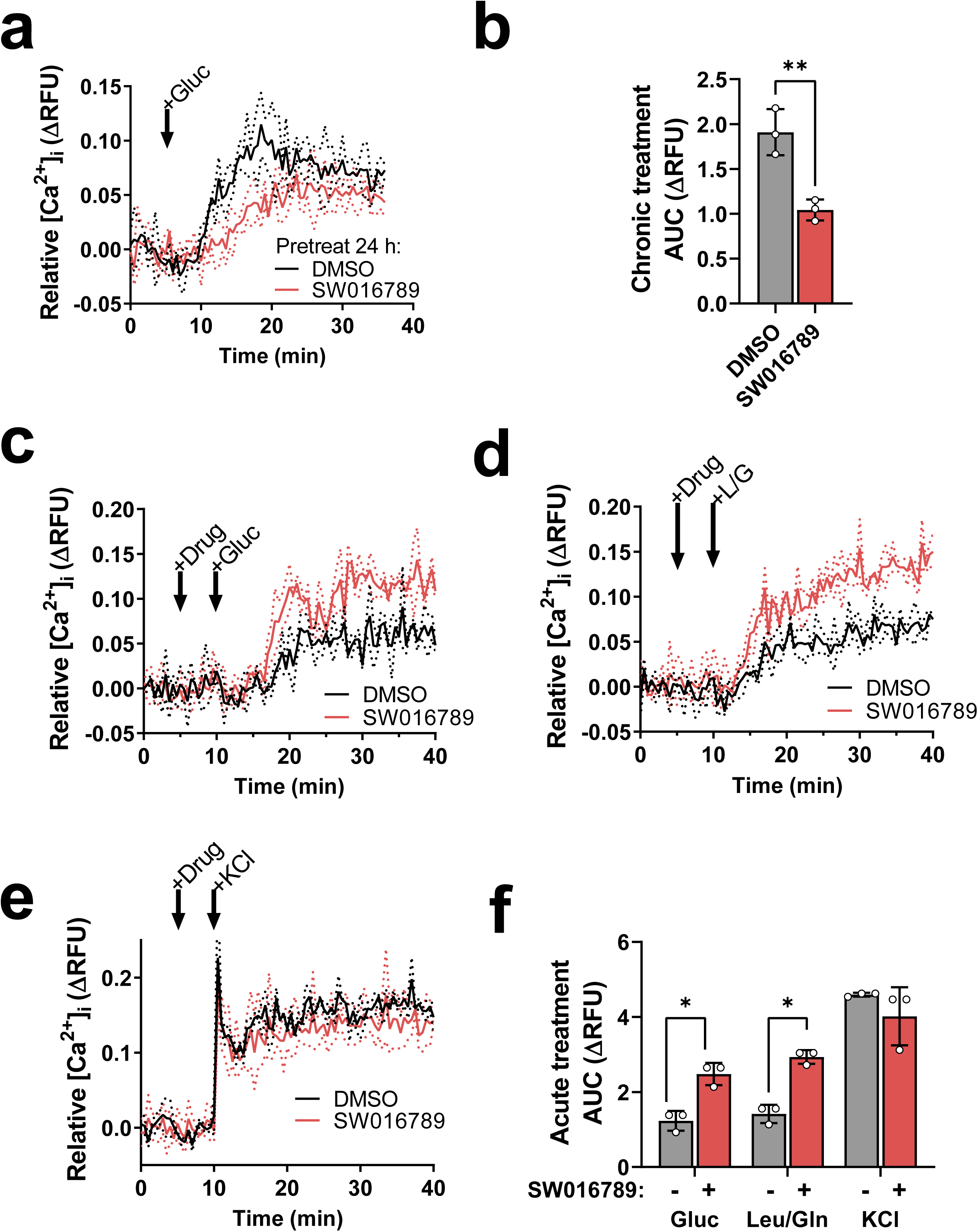
SW016789 treatment modulates nutrient-stimulated Ca^2+^ influx. **(a)** MIN6 beta cells pre-treated 24 h with SW016789 (5 µM) or DMSO (0.1%) followed by washout prior to glucose-stimulated Ca^2+^ influx assays and **(b)** area-under-the-curve (AUC) calculation. *, P<0.05 by unpaired Student’s t-test. **(c-f)** MIN6 beta cells acutely treated with SW016789 (5 µM) or DMSO (0.1%) followed by **(c)** glucose-, **(d)** leucine/glutamine-, or **(e)** KCl-stimulated Ca^2+^ influx assays and **(f)** associated AUCs. Data are the mean ± SD of three independent experiments. *, P<0.05 by two-way ANOVA.

**Figure 4.**
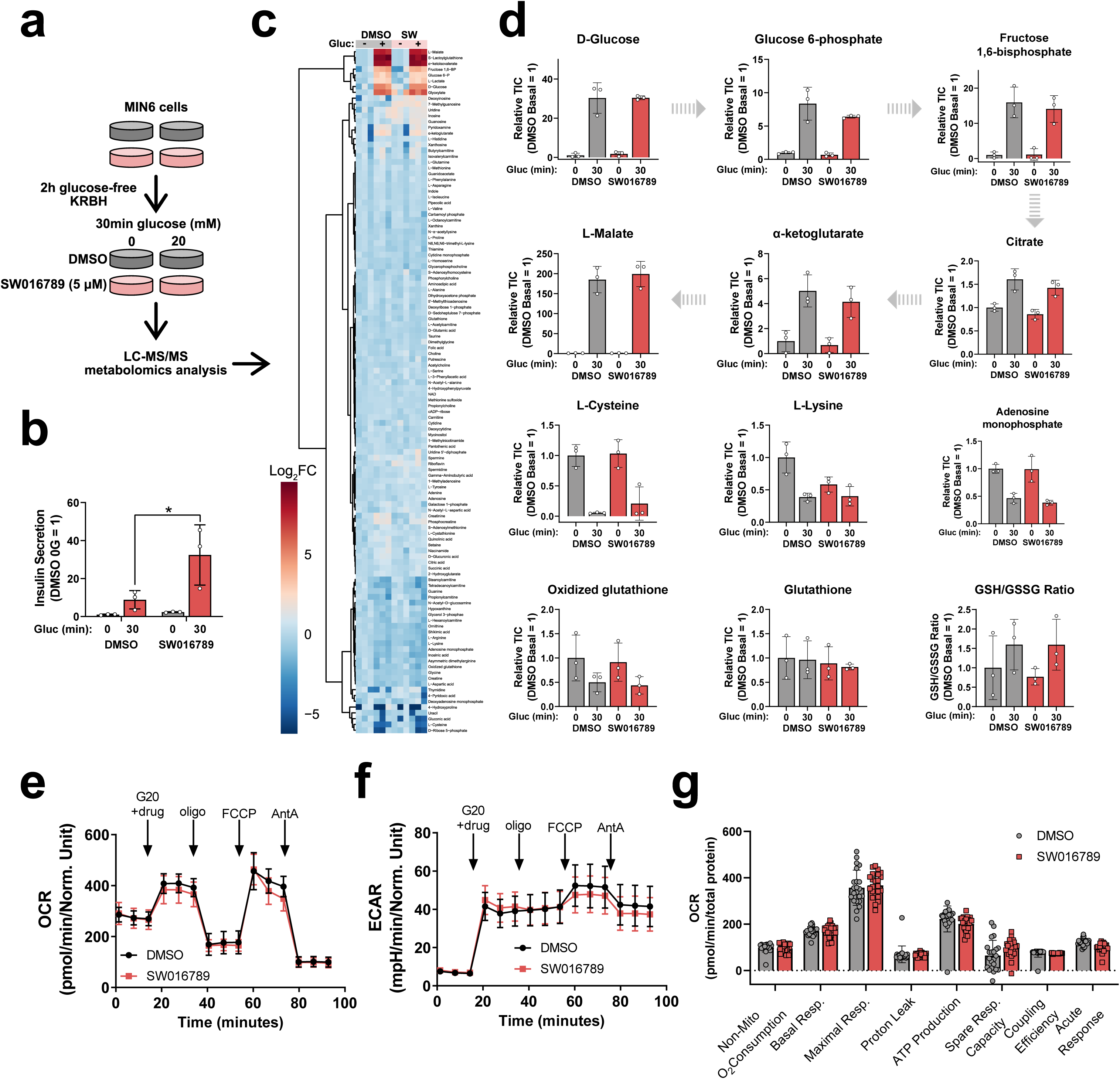
SW016789 does not impact oxygen consumption rate or canonical glycolysis and TCA cycle metabolites in MIN6 beta cells. **(a)** Experimental design for targeted metabolomics in MIN6 cells preincubated in glucose-free KRBH for 2 h and then treated with or without glucose (0 vs 20 mM) in the presence of DMSO (0.1%) (DMSO_0G and DMSO_20G) or SW016789 (5 µM) (SW_0G and SW_20G). **(b)** KRBH supernatants were collected for insulin ELISA to verify that SW016789 potentiated insulin secretion under these conditions. **(c)** Metabolomic data are represented as the log_2_ fold-change with respect to DMSO_0G of three independent experiments. **(d)** Representative metabolites showing glucose induced changes in glycolysis (glucose 6- phosphate, fructose-1,6-bisphosphate), TCA cycle (citrate, α-ketoglutarate, and malate), amino acids (cysteine, lysine), adenosine monophosphate, and redox potential (reduced glutathione (GSH) / oxidized glutathione (GSSG)) in MIN6 cells. **(e-g)** MIN6 beta cells were subjected to Seahorse analysis. Cells were preincubated in glucose-free KRBH and **(e)** oxygen consumption rate (OCR), **(f)** extracellular acidification rate (ECAR), and **(g)** calculated Seahorse parameters were measured during stimulation with glucose (20 mM) in the presence or absence of SW016789 (5 µM) or DMSO (0.1%). Subsequently all cells were sequentially stimulated with oligomycin, FCCP, and Antimycin A. Data represent the mean ± SD of three independent experiments.

### Chronic amplification of secretion leads to transient UPR^ER^ and shutdown of beta cell function

Acute SW016789-induced insulin hypersecretion eventually caused beta cells to lose normal GSCa and GSIS responses which we hypothesized was due to the UPR^ER^. To test this, we treated MIN6 cells in a time-course with SW016789 or thapsigargin and collected RNA and protein for analysis (**Fig. 5a**). Acute SW016789 increased immediate early gene (IEG) expression within 1 h which decreased to near baseline by 2-6 h (**Fig. 5b**), in agreement with observed GSCa effects. Thapsigargin did not affect expression these IEGs. SW016789 also induced expression of UPR^ER^ genes by ∼1 h with a peak from 2-6 h depending on the specific gene (**Fig. 5c**), followed by a return to near baseline by 24 h. This was in stark contrast to thapsigargin which caused a similar fold induction, but delayed peak of UPR^ER^ gene expression which persisted at 24 h.

**Figure 5.**
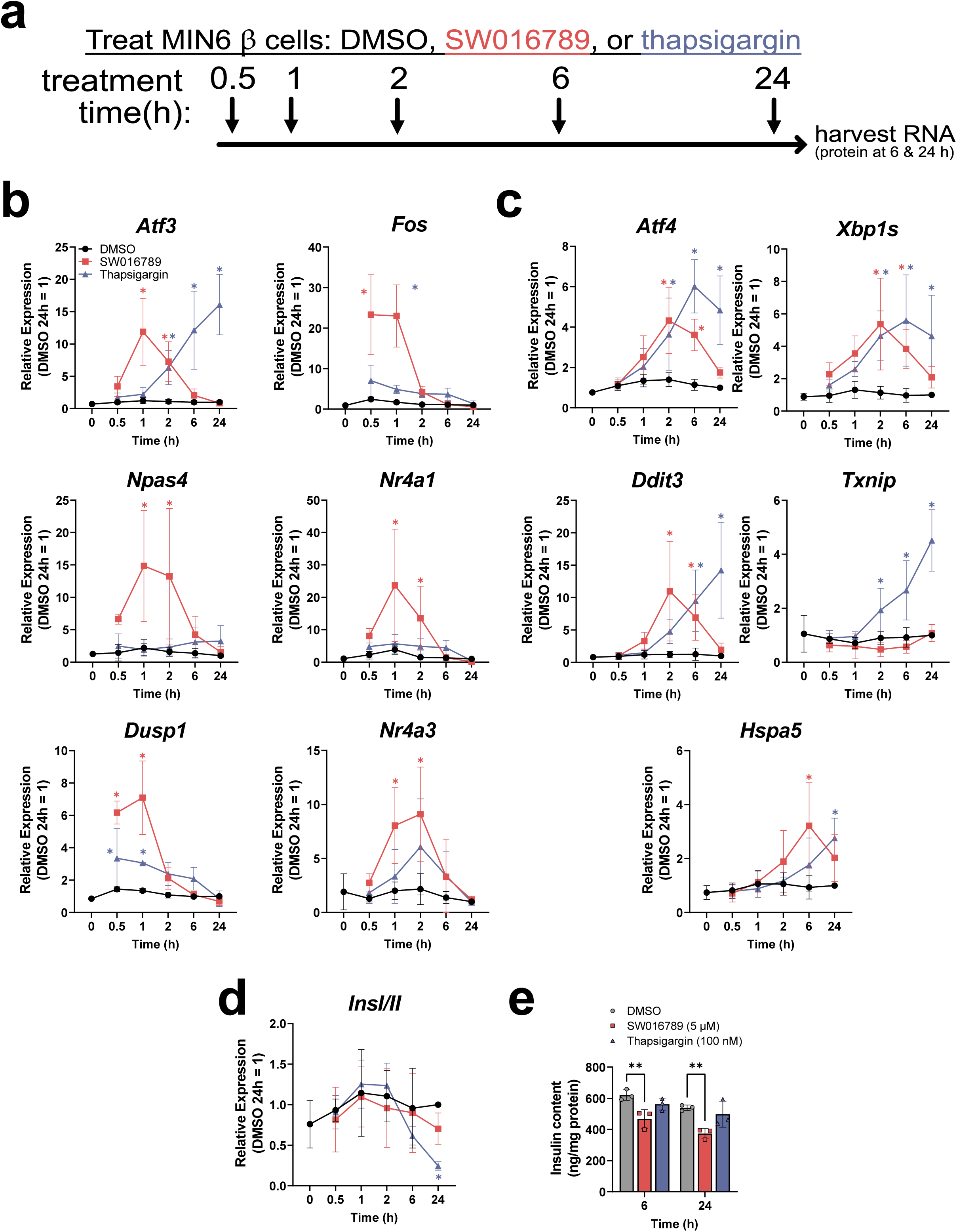
SW016789 transiently induces the unfolded protein response (UPR^ER^). **(a)** MIN6 beta cells were treated with DMSO (0.1%), SW016789 (5 µM), or thapsigargin (Tg, 100 nM) in complete medium for 0-24 has indicated. **(b-d)** Compound additions were staggered to allow simultaneous sample harvesting. Expression of **(b)** immediate early genes and **(c)** adaptive and maladaptive UPR^ER^ genes are shown. **(d)** Expression of *Ins1/2* in the time course and **(e)** insulin content. All data are the mean ± SD of three independent experiments. *, P<0.05 vs DMSO.

Notably, the oxidative stress transcription factor *Txnip* was induced only by thapsigargin. Over the time-course, SW016789 did not significantly alter insulin gene expression, but thapsigargin caused a decrease after 24 h (**Fig. 5d**). SW016789 decreased insulin protein content at 6 h but caused no further decrease by 24 h (**Fig. 5e**), suggesting the effect may be due to the induced hypersecretion and implied that after shutdown of beta cell function the drop in insulin content halted. We observed a similar pattern of induction and return to baseline of UPR^ER^ genes with higher doses of SW016789 (**ESM Fig. 4a**), indicating SW016789 did not induce terminal UPR^ER^ as occurs with thapsigargin. We also tested other endocrine cells that express stimulus-secretion coupling machinery similar to beta cells, including a human delta cell line and a mouse intestinal L cell line. SW016789 had minimal effects on IEGs or UPR^ER^ genes in these lines, while thapsigargin robustly induced the UPR^ER^ (**ESM Fig. 4b-c**).

We also observed differences at the protein level between SW016789 and thapsigargin for UPR^ER^ and apoptotic components (**Fig. 6a**). Thapsigargin, but not SW016789, significantly induced the apoptosis markers CHOP and cleaved PARP by 24 h. Phosphorylated PERK was significantly induced by thapsigargin at both 6 and 24 h, while the effects of SW01789 were minimal. Alterations in phosphorylated eIF2α were variable and not significantly different with any treatment at 6 and 24 h; however, thapsigargin induced BiP by 24 h. ATF3 and ATF4 protein expression was induced only at 6 h by SW016789 and to a lesser extent than with thapsigargin (**Fig. 6b**). Amounts of beta cell transcription factors Pdx1 and MafA were unaltered by 6 or 24 h of SW016789 or thapsigargin (**Fig. 6c**). Together, these results partially explain why even though SW016789 amplified secretion and caused ER stress, beta cells only ceased their secretory function and did not undergo terminal UPR^ER^.

**Figure 6.**
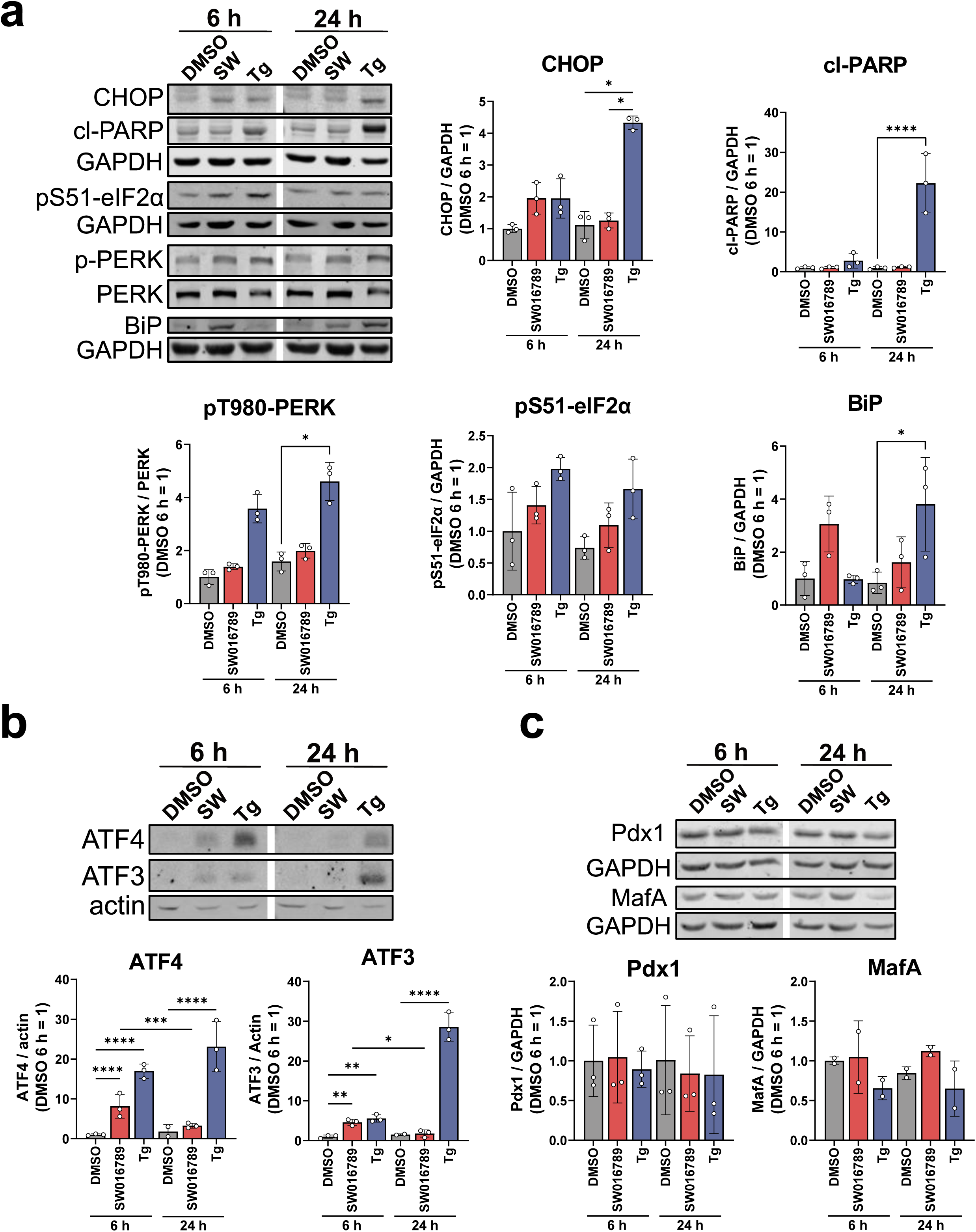
SW016789 and thapsigargin have distinct effects on UPR^ER^-related proteins. **(a)** MIN6 beta cells were treated with SW016789 (5 µM) or thapsigargin (Tg, 500 nM) for either 6 or 24 h in complete medium and lysates were immunoblotted. Thapsigargin, but not SW016789, induced pro-apoptotic markers CHOP and cleaved-PARP (cl-PARP). Alterations in UPR^ER^ markers pS51-eIF2α, pT980-PERK, and BiP are shown. **(b)** SW016789 and Tg increased the stress-induced transcription factors ATF4 and ATF3. Data are the mean ± SD of 3 independent experiments. **(c)** Beta cell transcription factors Pdx1 and MafA were not significantly affected by either drug. Data are the mean ± SD of 2-3 independent experiments. In all immunoblots either actin or GAPDH were used as loading controls. *, P<0.05 by two-way ANOVA.

### Acute effects of SW016789 are mediated via Ca^2+^ influx and PKCs

The UPR^ER^ effects of SW016789 in beta cells appeared to stem from the enhancement of nutrient-stimulated Ca^2+^ influx and insulin secretion. To elucidate the mechanisms underlying SW016789-induced hypersecretion of insulin and subsequent UPR^ER^ induction, we took two chemical approaches. In the first approach, we co-treated beta cells acutely with SW016789 and candidate compounds to identify any synergy or disruption in SW016789 actions. Among the stimuli tested, we discovered SW061789 exhibited synergy between and phorbol esters as well as Ca^2+^ channel openers. PKC activation by the phorbol ester phorbol 12-myristate 13-acetate (PMA) has well known effects on beta cells [34, 35]. Phorbol esters, including PMA, activate a subset of PKCs, as well as other targets [36], to potentiate nutrient-stimulated insulin secretion. While SW016789 and PMA each independently amplified GSIS, the combination of PMA and SW016789 elicited a marked synergistic insulin secretion response in the absence of nutrients (**Fig. 7a, DMSO pretreated**). To further explore the role of PMA-activated PKCs in the response to SW016789, we exploited the property of chronic (>16 h) treatment with PMA which causes the degradation of the PMA-responsive PKCs [35]. According to this paradigm, we pre-treated beta cells overnight with PMA and repeated the acute stimulation experiment. Under these conditions, SW016789 and PMA no longer stimulated secretion in the absence of nutrients and the effect of SW016789 on GSIS was no longer significant (**Fig. 7a, PMA pretreated**). These results indicated SW016789 may act in part though PKCs to elicit its acute effects. We also tested the effects of directly opening VDCCs in the presence or absence of SW016789. The potent VDCC activator FPL64176 phenocopied SW016789 in chronic (**Fig. 7b**) and acute (**Fig. 7c**) assays in beta cells. FPL64176 had distinct characteristics, including acutely synergizing with SW016789 in the absence of nutrients to induce Ca^2+^ influx and insulin secretion (**Fig. 7c,d**) and impacting GSCa almost immediately compared to the more delayed effect of SW016789 (**Fig. 7d**). These data suggested that FPL64176 and SW016789 act by different mechanisms.

**Figure 7.**
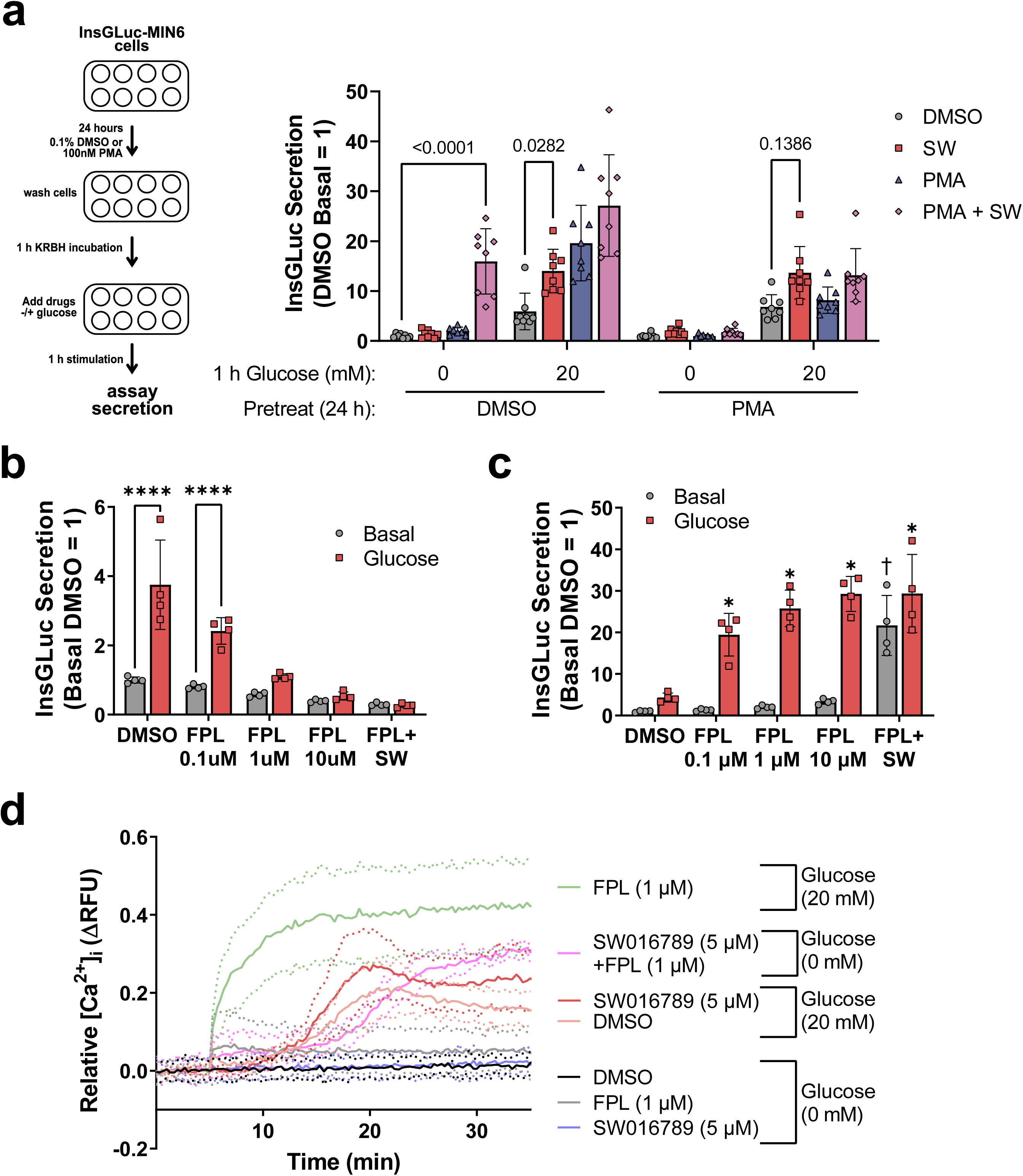
SW016789 acutely synergizes with activators of PKC and VDCCs to induce secretion. **(a)** InsGLuc-MIN6 cells were treated 24 h with either DMSO (0.1%) or PMA (100 nM) in complete medium. Cells were then washed and preincubated in glucose-free KBRH for 1 h prior to stimulation with SW016789 (5 µM), PMA (100 nM), or both in the presence or absence of glucose (20 mM) for 1 h. Afterward, buffer was collected for InsGLuc assays. Data are the mean ± SD of eight independent experiments. *,P<0.05 by two-way ANOVA. **(b)** InsGLuc-MIN6 cells were treated for 24 h with DMSO or the VDCC activator FPL64176 followed by glucose-stimulated InsGLuc secretion assay in the absence of compounds. Data are the mean ± SD of four independent experiments. *,P<0.05 Basal vs. Glucose. **(c)** InsGLuc-MIN6 cells were acutely treated for 1h with DMSO, FPL64176, or FPL64176 + SW016789 in the presence or absence of glucose. Data are the mean ± SD of four independent experiments. *,P<0.05 vs DMSO. **(d)** Normal GSCa can be observed with glucose and DMSO (peach) and is enhanced by SW016789 (red). FPL64176 rapidly potentiates glucose-stimulated Ca^2+^ influx in MIN6 cells (green). In the absence of glucose, DMSO (black), FPL64176 (grey) and SW016789 (blue) have little effect individually, but together FPL64176 and SW016789 synergize to stimulate Ca^2+^ influx (pink). Data are the mean ± SD of three independent experiments.

In a second approach we used a chemical suppressor screen to disrupt potentially involved pathways by co-treating cells with SW016789 and with compounds having known mechanisms. Compounds were tested for their ability to protect against chronic inhibitory effects of SW016789 (**Fig. 8a, ESM Table 2**). The VDCC blocker nifedipine was identified as a hit and was selected for further studies. Co-treatment of InsGLuc-MIN6 beta cells with SW016789 and nifedipine significantly blocked the chronic inhibitory effects of SW016789 on GSIS (**Fig. 8b**) and partially mitigated the induction of *Ddit3* (CHOP) expression at 6 h (**Fig. 8c**). We confirmed the acute potentiating effects of SW016789 on GSIS were indeed dependent on VDCC-mediated Ca^2+^ influx, as co-treatment with nifedipine completely blocked all stimulated secretion in the presence or absence of SW016789 (**Fig. 8d**). Furthermore, the protective effects of nifedipine against chronic SW016789 exposure were confirmed in human islet static culture GSIS experiments (**Fig. 8e**).

**Figure 8.**
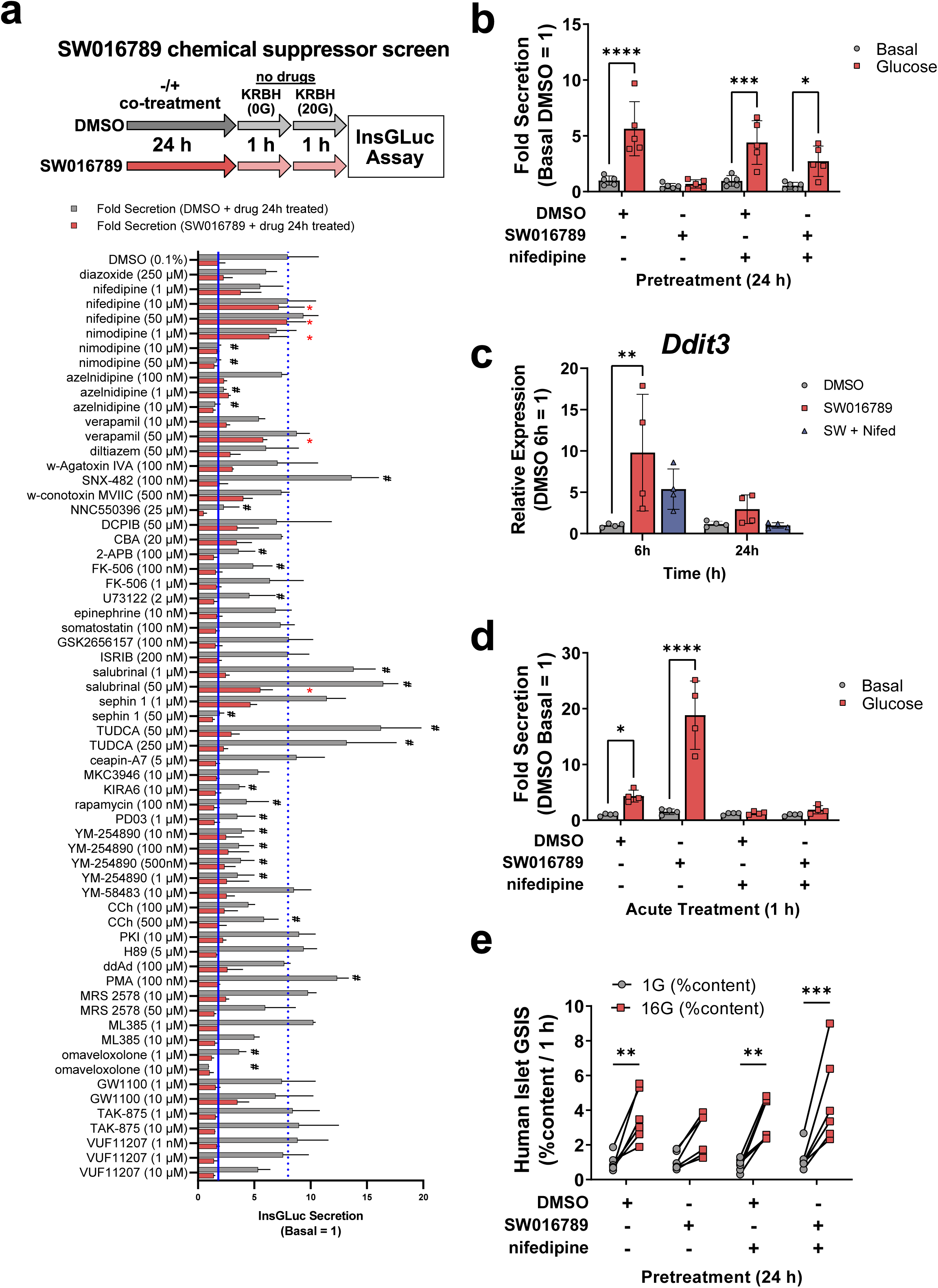
Chemical suppressor screen uncovers VDCC inhibitors as protective agents in chronic SW016789 treatment in MIN6 beta cells and human islets. (**a**) InsGLuc-MIN6 cells were co-incubated with SW016789 (5 µM), and candidate small molecules as indicated. Following 24 h incubation, all treatments were removed and glucose-stimulated InsGLuc secretion responses were assessed. (**b**) InsGLuc-MIN6 cells pretreated in complete medium for 24 h with DMSO (0.1%) or SW016789 (5 µM) in the presence or absence of nifedipine (10 µM). After compound washout cells were stimulated with or without glucose (20 mM) for 1 h for InsGLuc assays. Data are the mean ± SD of five independent experiments. (**c**) Cells were treated as in (b) and *Ddit3* gene expression was measured. Data are the mean ± SD of four independent experiments. (**d**) InsGLuc-MIN6 cells were acutely incubated with DMSO (0.1%), nifedipine (10µM), SW016789 (5µM), or nifedipine + SW016789 for 1h in the presence or absence of glucose (20mM). Data are the mean ± SD of four independent experiments. (**e**) Human islets were treated 24 h in complete medium with DMSO (0.1%) and SW016789 (5 µM) in the presence or absence of nifedipine (10 µM). Islets were washed and incubated in low glucose (1 mM) KRBH for 1 h, buffer discarded, then incubated 1 h in low (1 mM) or high (16 mM) glucose KRBH for 1 h. Data represent the mean ± SD of insulin measured in supernatant normalized to total insulin content from six different donor islet preparations. *, P<0.05 by two-way ANOVA.

## DISCUSSION

### Deconvolving the mechanism of action of SW016789

A few conclusions can be made about the mechanism of SW016789 based on the acute phenotype in beta cells. GSCa and GSIS were enhanced by SW016789, but KCl-induced responses were unaffected, suggesting the involvement of metabolism. Metabolic measurements suggested that SW016789 affects nutrient-induced insulin secretion downstream of ATP generation to enhance Ca^2+^ influx. We discovered intersections of SW016789-mediated actions with the PKC activator PMA and VDCC activator FPL64176. This raised concerns about the requirement for nutrient-stimulated ATP production given that SW016789, PMA, and FPL64176 are not nutrients and individually do not stimulate substantial Ca^2+^ influx or insulin secretion (**ESM Fig. 5**). In beta cells, phorbol esters potentiate GSIS through PKC activation [37, 38], but PKCs are not required for normal GSIS [35, 39]. Synergy between phorbol esters and SW016789 may occur through PKC-mediated phosphorylation of substrates involved in insulin exocytosis that lower the threshold for SW016789 to cause enhanced Ca^2+^ influx [40]. We also observed a small, but clear impact of SW016789 on Ca^2+^ influx and InsGLuc secretion in the presence of diazoxide and low KCl which was glucose-dependent (**Fig. 2c, ESM 2c)**. This suggests SW016789 can bypass K_ATP_ to cause membrane depolarization or VDCC opening. SW016789 could potentially outcompete the effect of diazoxide on K_ATP_, although we think this is unlikely because of estimations that >90% of K_ATP_ channels must close before beta cells depolarize [3, 41, 42].

Given the prominent role of cAMP in the potentiation of insulin secretion, we had initially suspected SW016789 might act through a Gαs-coupled GPCR. However, we did not observe a major impact of SW016789 in cAMP reporter assays (**ESM Fig. 3a-b**). We also did not observe protection from chronic SW016789 effects by co-treatment with inhibitors of cAMP pathways (e.g. PKA inhibitor H89, adenylyl cyclase inhibitor ddAd) (**Fig. 8A**). While previous work has shown that the cAMP-PKA pathway can protect against thapsigargin-mediated beta cell death [43], SW016789 did not cause cell death (**Fig. 1g-i**) and the UPR^ER^ dynamics induced by SW016789 appear to be quite different compared to thapsigargin.

Among compounds tested for protection against SW016789, the most consistent hits were from the dihydropyridine VDCC inhibitor family. We surmise that preventing increased Ca^2+^ influx abrogates SW016789-enhanced insulin secretion, thereby preventing downstream UPR^ER^ and preserving beta cell function. As blocking VDCCs prevents both Ca^2+^ influx and subsequent insulin secretion, it is unclear whether downstream stress and functional loss is induced specifically because of chronically increased Ca^2+^, enhanced insulin release, or both. Indeed, genetically reducing insulin production by 50% mitigated stress in mouse beta cells [44], suggesting the possibility that in our system, blocking insulin secretion itself using VDCC inhibitors may be responsible for mitigating hypersecretion-induced stress.

### Beta cell adaptation and dysfunction in response to chronic secretory demand

Chronic metabolic stress leads to beta cell dysfunction and two recent studies provide prime examples [45, 46]. Sustained physiological hyperglycemia (72 h) increases absolute insulin secretion but impairs beta cell function as gauged by the disposition index [45]. Accordingly, human islets exposed to glucolipotoxic stress for 48 h were unable to recover GSIS, while islets exposed to milder stress (high glucose or palmitate alone) were able to recover function [46].

Under our culture conditions SW016789 initially potentiated beta cell function, but the secretory response shut down by 4 h. Although UPR^ER^ induction was transient, returning to baseline after 24 h, beta cell function in terms of insulin secretion remained impaired. Beta cells recovered from this inhibited state after a 48 h washout, indicating the changes were reversible. A potential explanation is that a protein(s) affected by SW016789 signaling are degraded/reduced and require the washout period to recover. Recently, multiple groups have shown that transient UPR^ER^, either downstream of partial pancreatectomy [47] or via inhibition of salt-inducible kinase (SIK) [48], is associated with enhanced beta cell proliferation. While we did not observe significant alterations in cell viability or death over a 72 h exposure to SW016789, it is possible the mechanism of UPR^ER^ induction due to hypersecretion in this case is different from that of pancreatectomy or SIK inhibition.

SW016789 selectively and transiently induced canonical UPR^ER^ stress response genes, but with a distinct temporal pattern from that of thapsigargin. A lesser induction of ATF3/4, p-PERK, and p-eIF2α may indicate an overall weaker induction of the PERK arm of the UPR^ER^ by SW016789. Our findings also suggest a lack of oxidative stress induction by SW016789 ascertained by no observed changes in glutathione redox state or *Txnip* expression. SW016789- treated beta cells were therefore able to downregulate their response to transiently enhanced Ca^2+^ influx. Relative induction of pro-survival versus pro-apoptotic genes may underlie the effects of SW016789-mediated UPR^ER^. For example, the Ca^2+^-activated gene *Npas4* was shown to have a protective role in beta cells and was highly induced by SW016789, but not by thapsigargin [49, 50]. A gene program that mitigates stress within the first 6 h of drug treatment may allow SW016789-treated beta cells to downregulate UPR^ER^ pathway before the point of no return. SW016789 will be a useful tool compound for discovering genes involved in adaptive versus maladaptive responses to chronic Ca^2+^ influx or secretory pressure.

### Role of pharmacological modulation of VDCC activity for beta cell protective effects

Ca^2+^ channel blockers have been in the spotlight for their therapeutic value in both type 1 [51] and type 2 diabetes [52]. For example, verapamil improved beta cell function in a clinical trial of recent-onset type 1 diabetes [51]. The proposed mechanism of action is that suppressing beta cell Ca^2+^ influx reduces expression of the cellular redox regulator *Txnip*, leading to protection from apoptosis [53]. In our studies SW016789 did not induce *Txnip*, suggesting an alternate mechanism. In addition, the VDCC blockers of the dihydropyridine class (nifedipine, nimodipine, amlodipine), as well as the phenylalkylamine class (verapamil) protected beta cells from SW016789-induced dysfunction. Our results showing the protective effects of nifedipine agree with previous findings in beta cells using other stressors including hyperglycemia and palmitate [54–56]. Distinct VDCC inhibitor mechanisms of action may underlie the differences between dihydropyridine and phenylalkylamine-mediated beta cell protection [57] and may imply that SW016789 acts through a mechanism or pathway which is more dependent on VDCCs in the inactivated state. Because amplified GSCa was confirmed as a major mechanism underlying the effects of SW016789, we hypothesized that directly activating VDCCs using FPL64176 could phenocopy SW016789 [58]. While SW016789 activity on Ca^2+^ influx was inhibited by nifedipine, it also synergized with FPL64176 to enhance Ca^2+^ influx and secretion. The kinetics of Ca^2+^ influx induced by FPL64176 or SW016789 in the presence of glucose differed greatly. In agreement with our findings with FPL64176 and SW016789, prior work showed that chronic treatment with VDCC activator BayK8644 caused insulin secretion defects attributed to decreased abundance of K_ATP_ channel and VDCC subunits [59]. Although not assessed in those studies, beta cells likely underwent transient UPR^ER^ concomitant with or prior to losing function.

Nifedipine or newer generation dihydropyridines like amlodipine, may also be useful therapeutics for conditions that cause hyperinsulinemia or where beta cell VDCCs are dysfunctional. A recent example is an individual with congenital hyperinsulinism with a gain-of-function mutation at a conserved residue in the α1 pore-forming subunit of Cav1.3 (L271H), encoded by *CACNA1D* [60]. Hyperinsulinism in this individual was successfully treated with nifedipine and diazoxide. However, chronic blockade of beta cell VDCCs should be carefully considered, as preliminary findings with genetic deletion of Cav1.3 in mice suggest a positive role in beta cell mass [61]. Auxiliary channel subunits can also impact beta cell function in diabetes. For example, the VDCC β_3_ subunit (*Cacnb3*) is upregulated in islets from diabetic animals and was associated with decreased channel function. Deletion of *Cacnb3* improved beta cell function in a mouse model of diabetes [62, 63]. Future research into precision targeting of VDCCs or their regulatory partners may be a promising avenue for diseases of beta cell dysfunction.

In conclusion, we discovered a new compound and potentially novel pathway that modulates beta cell function and may expand the cohort of targets for disease mitigation. Simultaneously, these discoveries also generate new knowledge about the mechanics of nutrient- and hormone-regulated exocytosis from specialized secretory cell types, including insights into the metabolic amplifying pathway.

## Acknowledgements

We thank past and present members of the Cobb lab, Whitehurst lab and D. Rosenbaum at UT Southwestern; D. Eizirik and M. Cnop at Université libre de Bruxelles; and A. Templin at IBRI for helpful discussions. Thanks to C. Cheng in the Whitehurst lab and C. Llamas in the Mishra lab at UT Southwestern for training and advice on Seahorse experiments. Thanks to the L. Zacharias and the DeBerardinis lab at UT Southwestern for advice on metabolomics analyses. Agonist and antagonist functional data on GPCRs for SW016789 were generously provided by B. Roth’s group through the National Institute of Mental Health’s Psychoactive Drug Screening Program, Contract # HHSN-271-2018-00023-C (NIMH PDSP). MIN6 beta cells were originally generated by J Miyazaki [64] and were kindly shared with us via the late J.C. Hutton. Human QGP1 somatostatinoma cells were a gift from D. Quelle (University of Iowa Carver College of Medicine). Mouse intestinal GLUTag L cells were originally generated by D. Drucker (Mount Sinai Hospital) and were shared with us by the lab of G.G. Holz (Upstate Medical University).

## Data availability

All data generated or analyzed during this study are included in this published article (and its electronic supplementary materials).

## Contribution statement

Conceptualization: M.A. Kalwat.

Data curation: M.A. Kalwat.

Formal analysis: K. Rodrigues-dos-Santos, G. Roy, D.D. Binns, F. Armoo, and M.A. Kalwat.

Funding acquisition: M.A. Kalwat and M.H. Cobb.

Investigation: K. Rodrigues-dos-Santos, G. Roy, D.D. Binns, M.G. Grzemska, F. Armoo, M.K. McCoy, A. Huynh, J.Z. Yang, and M.A. Kalwat.

Methodology: M.A. Kalwat.

Project administration: M.A. Kalwat.

Resources: B.A. Posner, M.H. Cobb, and M.A. Kalwat

Supervision: M.H. Cobb and M.A. Kalwat

Validation: K. Rodrigues-dos-Santos, G. Roy, D.D. Binns, M.G. Grzemska, J.Z. Yang, and M.A. Kalwat.

Visualization: K. Rodrigues-dos-Santos, G. Roy, and M.A. Kalwat.

Writing - original draft: M.A. Kalwat.

Writing - review & editing: K. Rodrigues-dos-Santos, G. Roy, D.D. Binns, L. Barella, M.H. Cobb, and M.A. Kalwat

M.A. Kalwat is the guarantor and accepts full responsibility for the work and/or the conduct of the study, had access to the data, and controlled the decision to publish.

## ESM Methods

**Electronic Supplementary Material (ESM):** Rodrigues-dos-Santos K, Roy G, Binns DD, et al.

### Ca^2+^ measurements in the diazoxide paradigm or for store-operated Ca^2+^ entry (SOCE)

For Ca2+ measurements in the presence of diazoxide, after loading with Ca2+ indicator dye, cells were incubated in KRBH containing 250 µM diazoxide for 5 min prior to the baseline read and subsequent stimulation with 35 mM KCl and/or 20 mM glucose with DMSO or SW016789 in the continued presence of diazoxide. For measurements of ER Ca2+ and SOCE, beta cells were incubated in normal glucose-free KRBH for 1 h, then loaded with Fura-2-LR-AM (5 µM) in Ca2+-free KRBH (5 mM KCl, 120 mM NaCl, 15 mM HEPES, pH 7.4, 24 mM NaHCO3, 3 mM MgCl2, and 1 mg/ml radioimmunoassay-grade BSA) for 25 min and a 5 min washout in the same buffer. Cells were placed in the BioTek Synergy H1 for a baseline read (30 s), followed by injection of thapsigargin (10 µM) for a 5 min read, and CaCl2 (2.5 mM) for another 5 min read.

### cAMP reporter assays

For relative cAMP measurements, the genetically-encoded Epac-based bioluminescence-resonance energy transfer (BRET) reporter for cAMP (CAMYEL) was used as previously described [1]. CAMYEL MIN6 cells were preincubated in glucose-free KRBH for 1.5 h. Cells were incubated for 15 min at room temperature with coelenterazine-h (20 µM). After a 3 min baseline recording, stimulations were added, and plate read for an additional 15 min. Using the BRET1 cube from BioTek, emission signals at 460/40 and 540/25 nm (center/bandpass) were measured every 0.8 s. Data were normalized by subtracting the mean of the respective baseline reads. Area under the curves (AUCs) were calculated in GraphPad Prism using a baseline of 0 ΔRFU and ignoring peaks that are <10% of the distance from minimum to maximum ΔRFU and those defined by fewer than 15 adjacent points. All peaks must go above the baseline.

### Chiral separation and structure-activity analysis

Separated enantiomers of racemic SW016789 were obtained from Lotus Separations using supercritical fluid chromatography [2]. Purified enantiomers were resuspended in DMSO and subjected to acute and chronic InsGLuc-MIN6 experiments as described. For structure-activity analysis, structures of SW016789 analogs and their respective activity scores were analyzed in Datawarrior software [3]. Glucose-stimulated InsGLuc secretory activity normalized to unstimulated secretion was input along with structures and activity-cliff analysis was performed.

## ESM Results

**ESM Figure 1.**
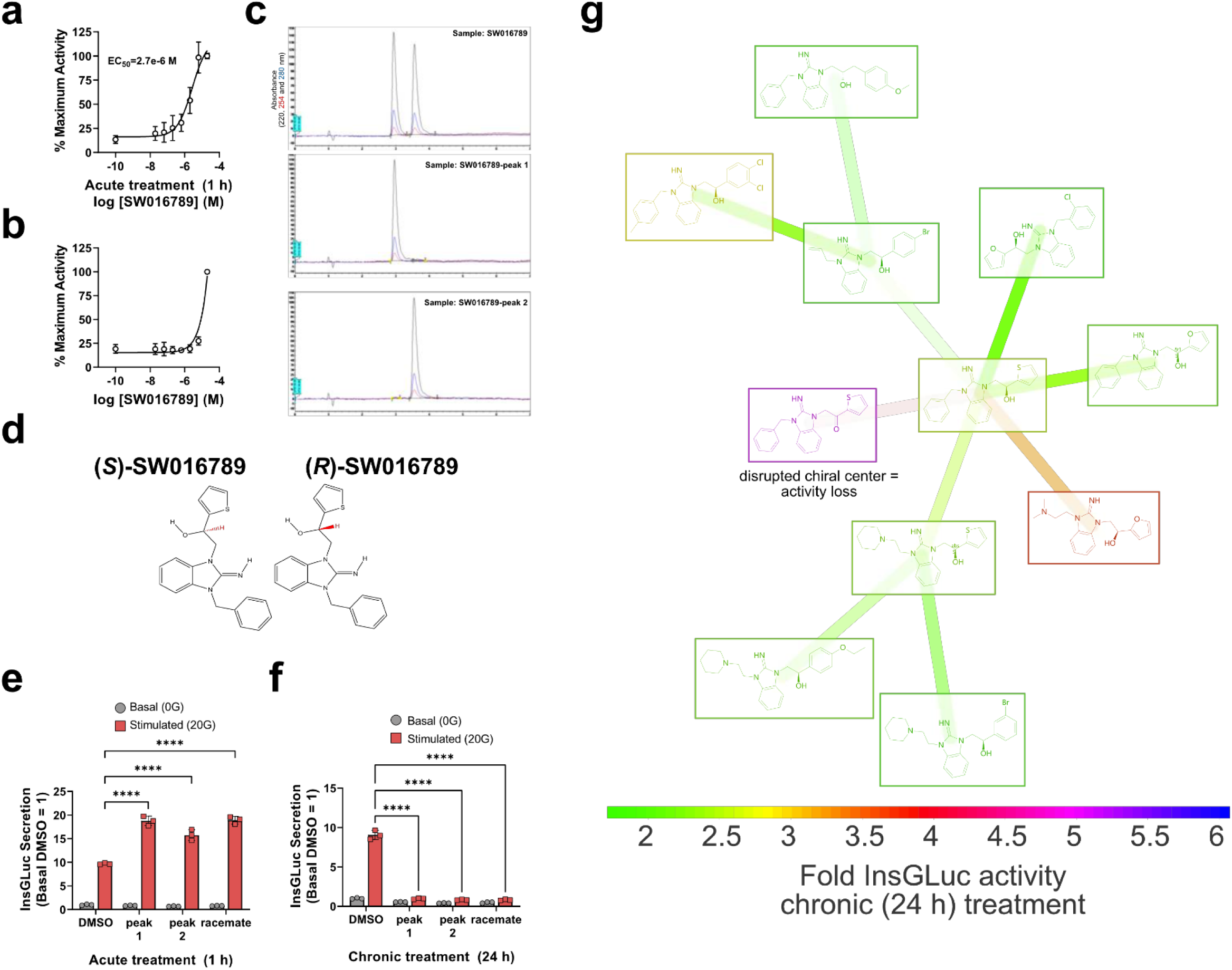
Acute dose-response testing of SW016789 in InsGLuc-MIN6 cells. Cells were treated with varying concentrations of SW016789 or vehicle (0.1% DMSO) in the **(a)** presence and **(b)** absence of glucose (20 mM) for 1 h. Data are the mean ± SD of three independent experiments. The EC_50_ of SW016789 calculated in the presence of glucose was 2.7E-6 M. **(c)** Chiral separation of racemic mixture of SW016789 into two distinct peaks. **(d)** Enantiomeric structures of SW016789. **(e,f)** Secreted Gaussia activity from InsGLuc-MIN6 cells treated either **(e)** only during glucose stimulation (1 h) or **(f)** chronically (24 h) prior to glucose stimulation. Cells were exposed to DMSO (0.1%), or purified enantiomers of SW016789 (peak 1, peak 2, 5 µM each), or the SW016789 racemate (5 µM). Data are the mean ± SD of three independent experiments. *, P<0.05 by two-way ANOVA. **(g)** Structure-activity relationship of structural analogs assayed in chronic (24 h) treatment of InsGLuc- MIN6 cells. Fold InsGLuc activity is normalized to secretion in unstimulated conditions. Increased fold activity indicates a loss in chronic inhibitory activity, as seen for the structure with disrupted chiral center. DataWarrior was used to generate a network display organized by structural similarity and colored by activity.

**ESM Figure 2.**
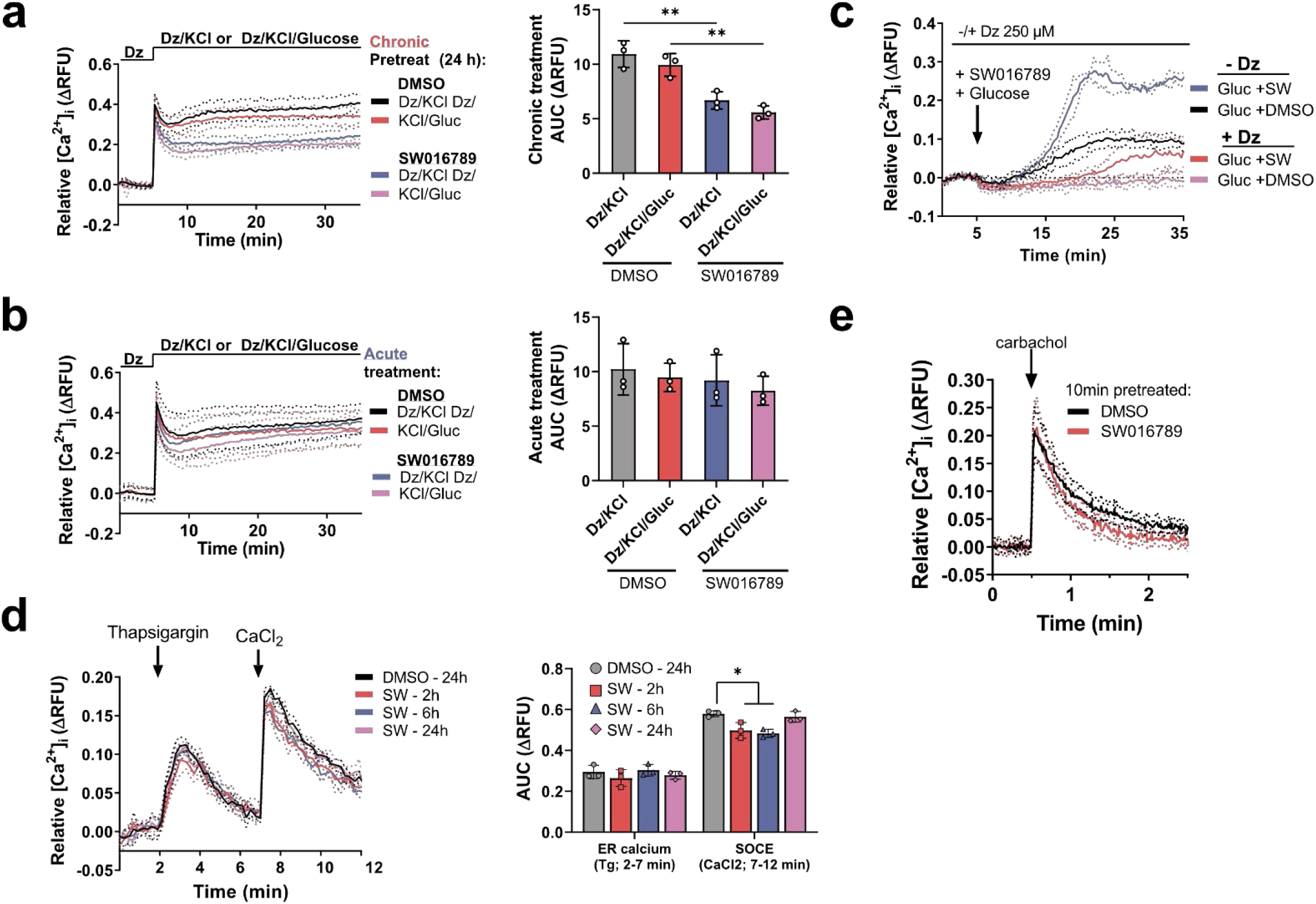
Effects of SW016789 on Ca^2+^ influx under the diazoxide paradigm and during SOCE and ROCE. **(a,b)** InsGLuc-MIN6 cells were exposed to DMSO (0.1%) or SW016789 (5 µM) for **(a)** 24 h before Ca^2+^ influx assay or **(b)** were acutely co-stimulated with DMSO or SW016789, as indicated above each graph. Cells were incubated in the presence of diazoxide (Dz, 250 µM) and stimulated with either Dz + KCl 35 mM or Dz + KCl 35 mM + glucose 20 mM. Expectedly, the addition of glucose did not alter the amount of Ca^2+^ influx in the presence of diazoxide and KCl in either chronic or acute treatments. Associated AUC calculations are shown. Data are the mean ± SD of three independent experiments. *,P<0.05 by one-way ANOVA. **(c)** MIN6 cells were incubated in the presence or absence of diazoxide (250 µM) in the KRBH during Ca^2+^-influx assays. Cells were then stimulated with glucose (20 mM), SW016789 (5 µM), or both at the time indicated by the arrow. Data are the mean ± SD of three independent experiments. **(d)** Endoplasmic reticulum (ER) Ca^2+^ stores and store-operated Ca^2+^ influx (SOCE) were assayed in MIN6 beta cells. Thapsigargin (Tg, 10 µM) elicited ER Ca^2+^ leakage into the cytosol and adding back CaCl_2_ (2 mM) resulted in SOCE. AUCs were calculated separately for each peak. **(e)** MIN6 cells were stimulated with carbachol (CCh, 250 µM) to assess receptor-operated Ca^2+^ entry (ROCE) after a 10 min pretreatment with SW016789 or DMSO. Data are the mean ± SD of three independent experiments.

**ESM Figure 3.**
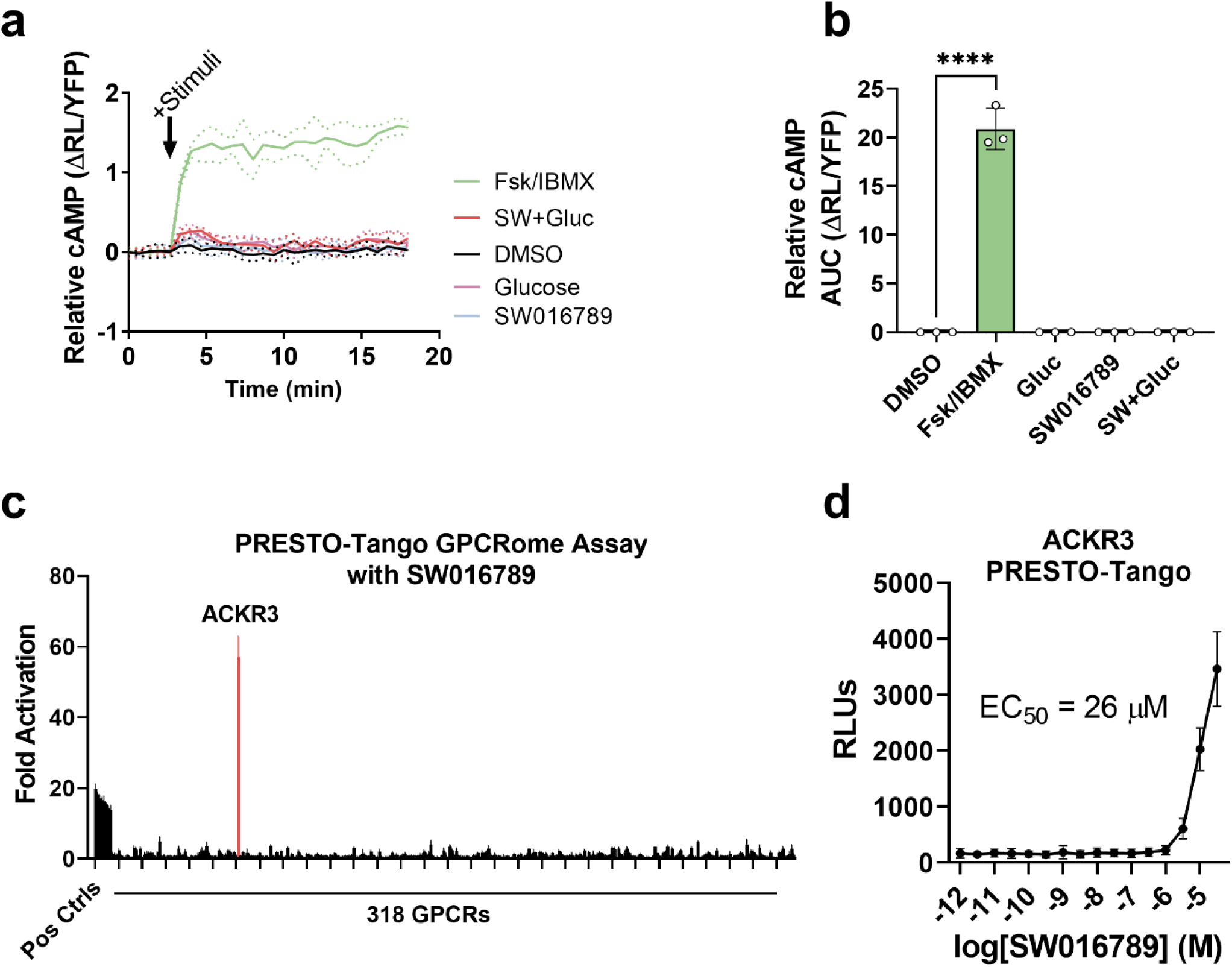
GPCR screening of SW016789 in the PRESTO-TANGO assay. **(a)** MIN6 cells stably expressing the cAMP BRET sensor CAMYEL were preincubated in glucose-free KRBH for 1.5 h prior to treating with coelenterazine-400A for 15 min. Baseline BRET was read for 3 min prior to stimulation as indicated. Cells were stimulated with forskolin (Fsk, 10µM) and IBMX (1µM) as a positive control, SW016789 (10 µM) alone, glucose (20 mM) alone, SW016789 and glucose together, or DMSO (0.1%). **(b)** ΔBRET peak area defined using baseline of 0 and ignoring peaks under 0.5 units high and fewer than 10 adjacent points. Data are the mean ± SD of 3 independent experiments as indicated by individual data points on the AUC bar graph. *, P<0.05 by two-way ANOVA. **(c)** Results of PRESTO-TANGO assay using 10 µM SW016789 showed only one prominent hit, CXCR7. **(d)** Validation studies performed at the PDSP core show the EC50 of SW016789 on CXCR7 is ∼26 µM.

**ESM Figure 4.**
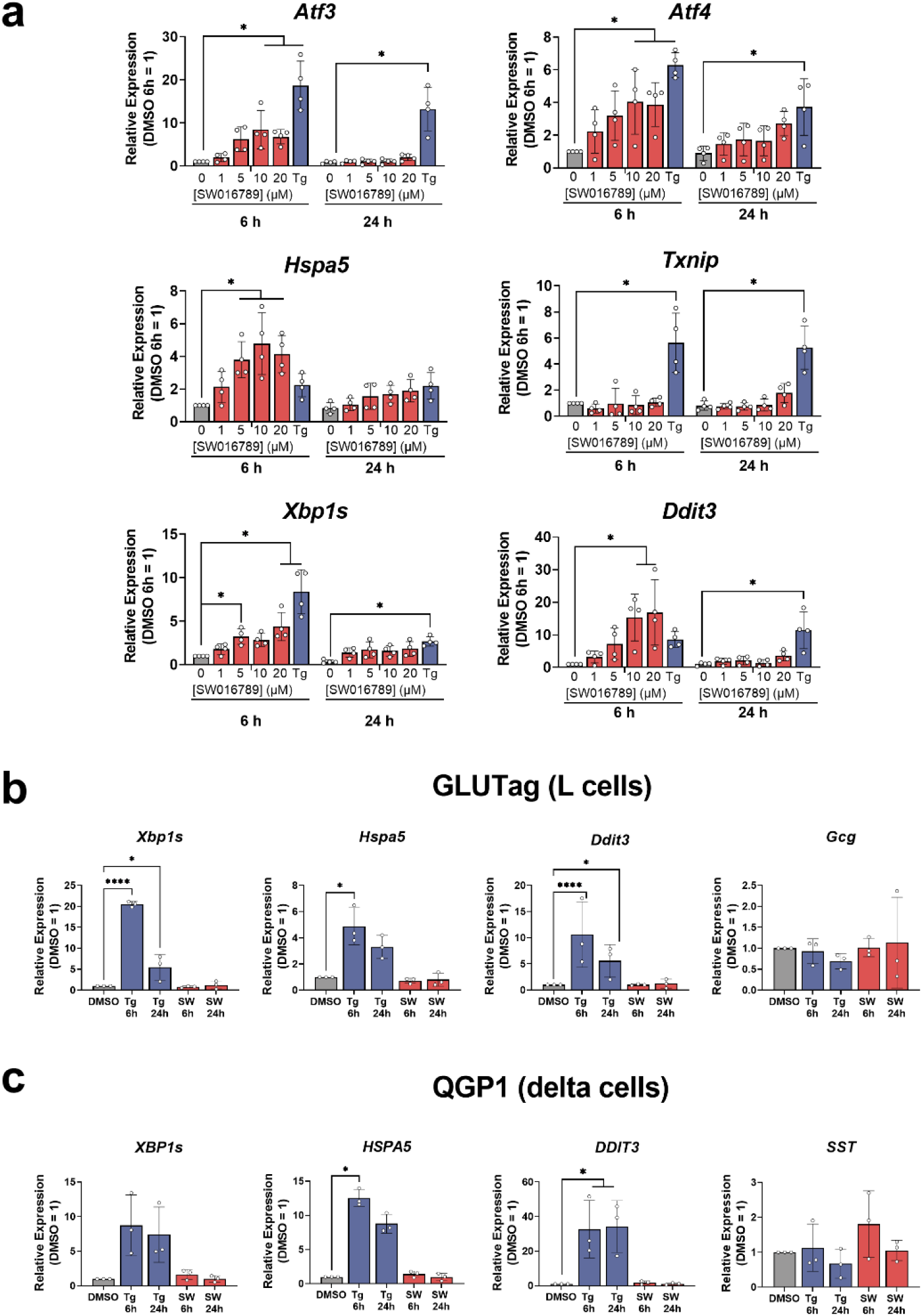
SW016789 did not impact UPR^ER^ in other neuroendocrine cells nor cause maladaptive UPRER at high concentrations. **(a)** Relative UPR^ER^ gene expression in MIN6 cells treated for either 6 or 24 h with vehicle (0.1% DMSO) or increasing concentrations of SW016789 (1-20 µM). Thapsigargin (100 nM) was included as a positive control at each time point. Data are the mean ± SD of four independent experiments. *, P<0.05 by two-way ANOVA. **(b,c)** Relative gene expression in a mouse intestinal L-cell cell line (GLUTag) **(b)** and a human islet delta cell line (QGP1) **(c)** treated with DMSO (0.1%), SW016789 (5 µM), or thapsigargin (Tg, 100 nM) in complete medium for 6 or 24 h. Glucagon (*Gcg*) message was measured for L-cells as they produce GLP-1 encoded by *Gcg*. Somatostatin (*SST*) was measured as the major product of δ cells. All data are the mean ± SD of at least 3 independent experiments. *,P<0.05 by two-way ANOVA.

**Table S1.**
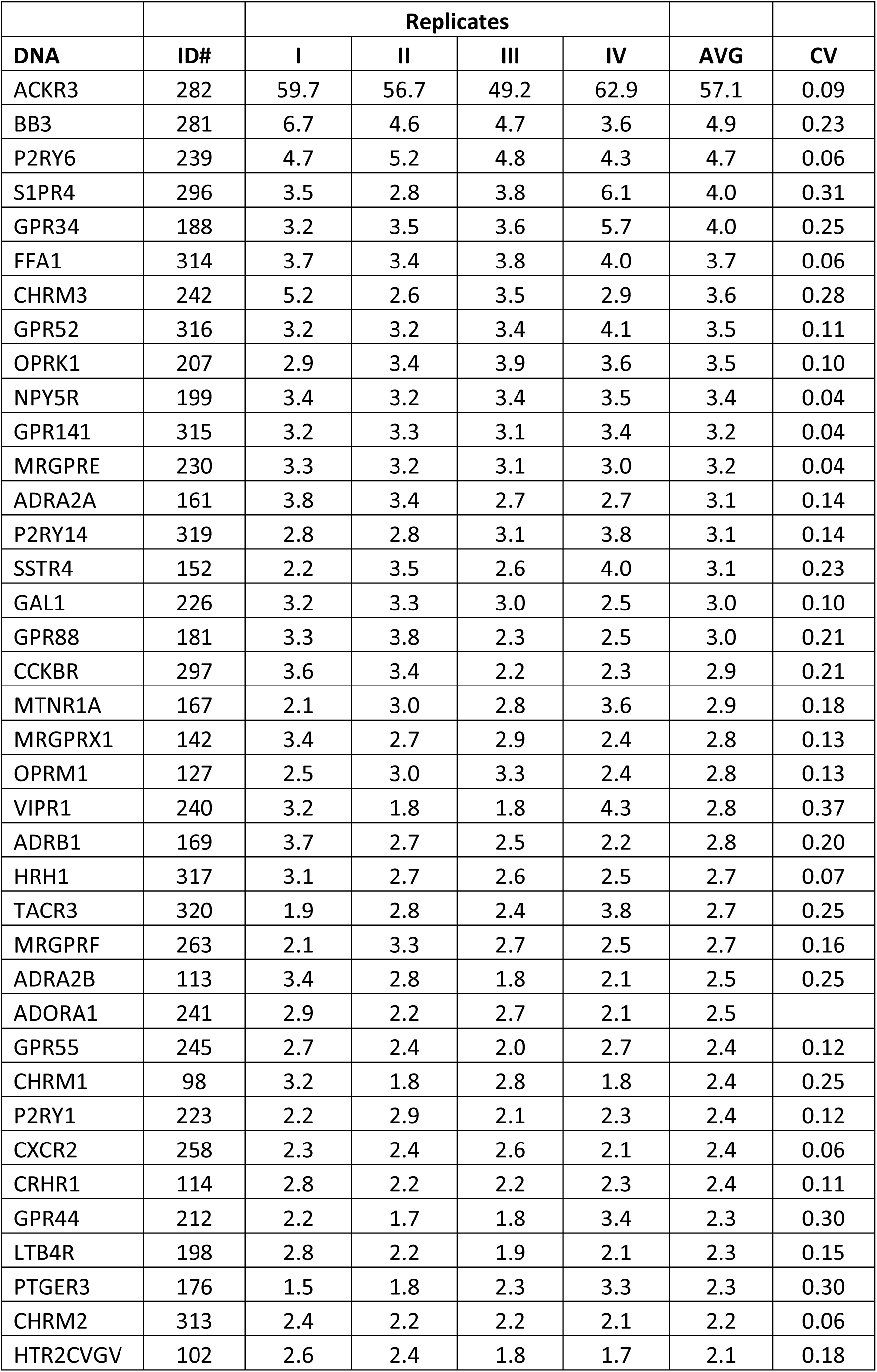
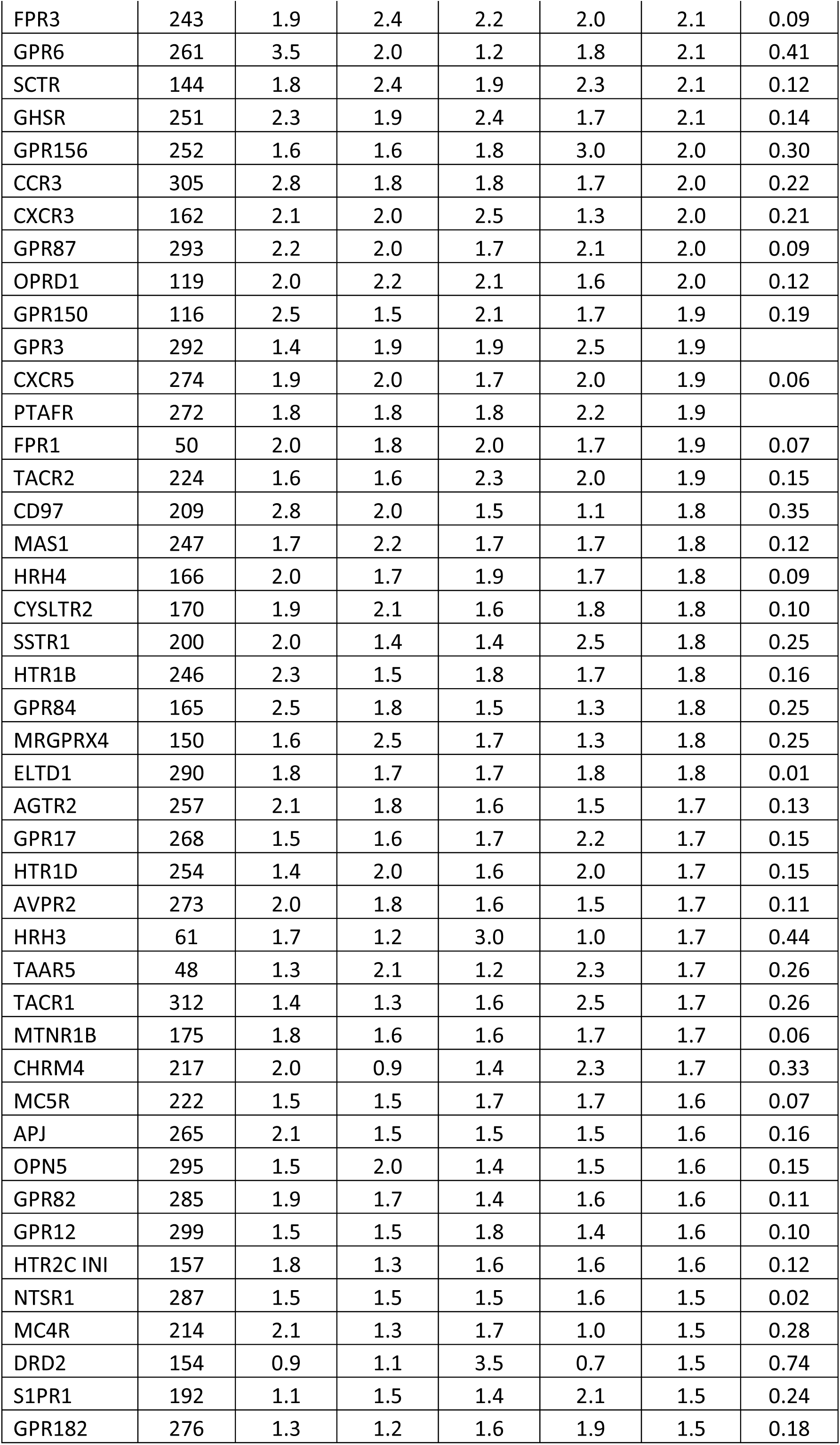
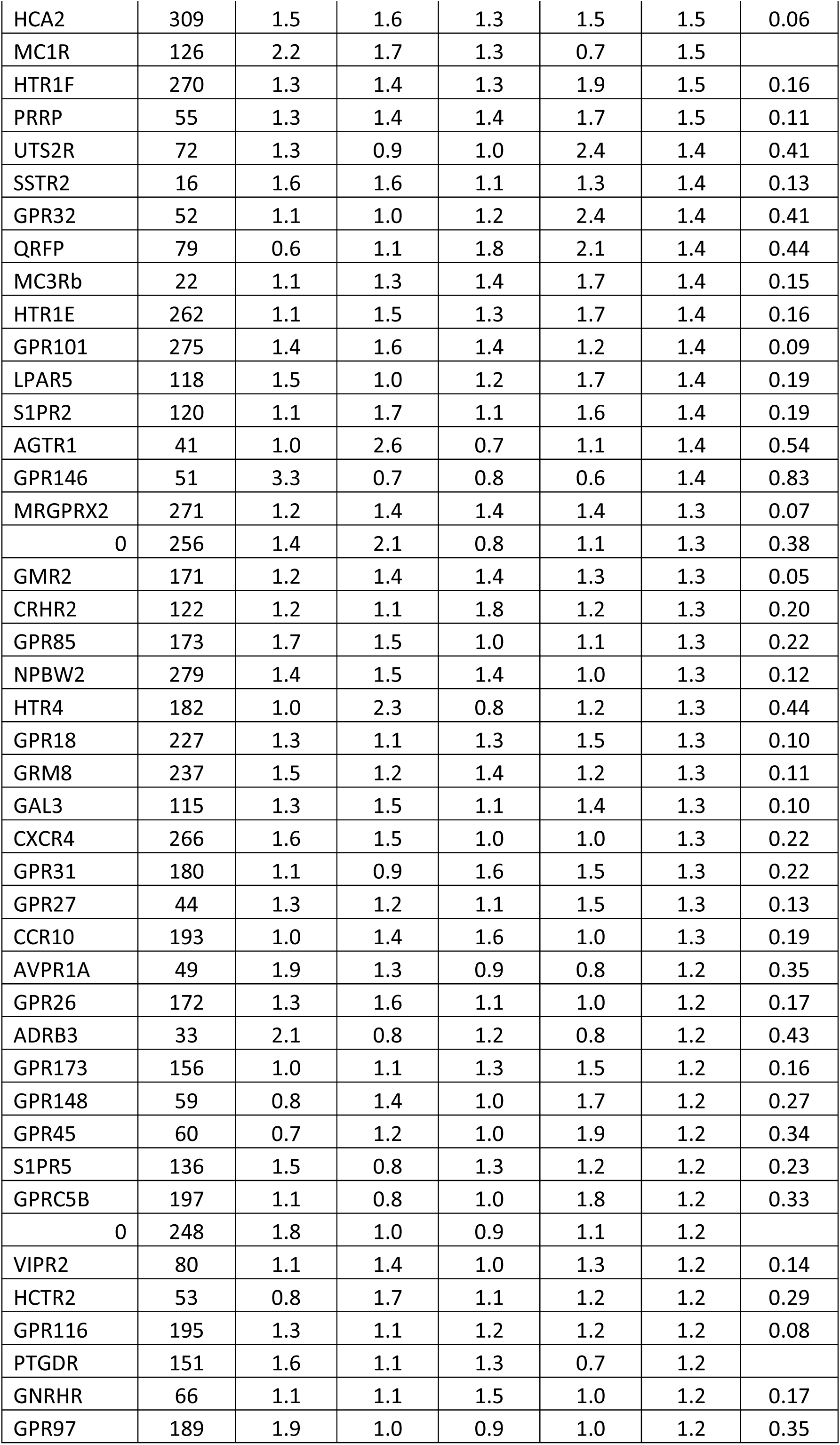
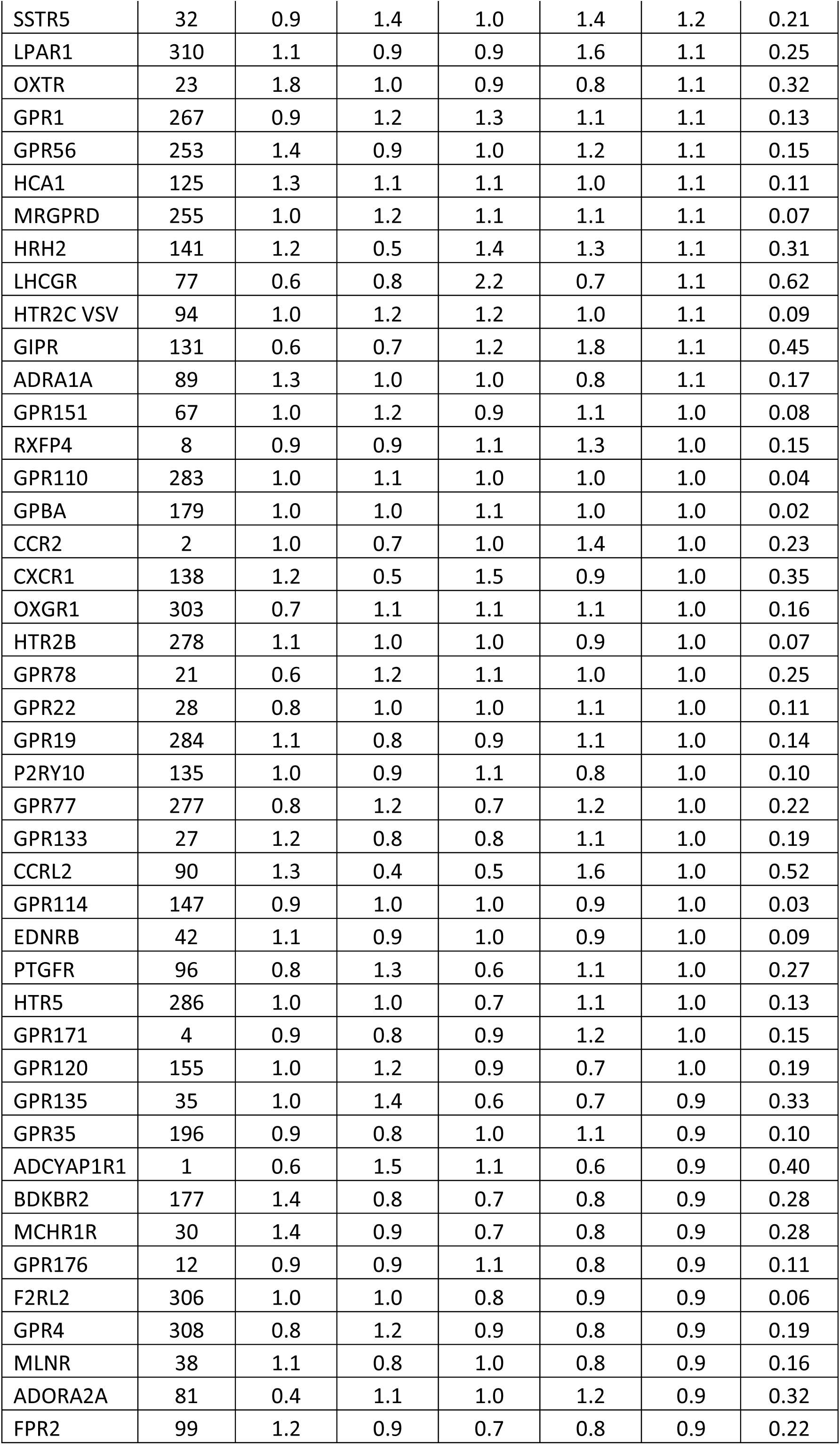
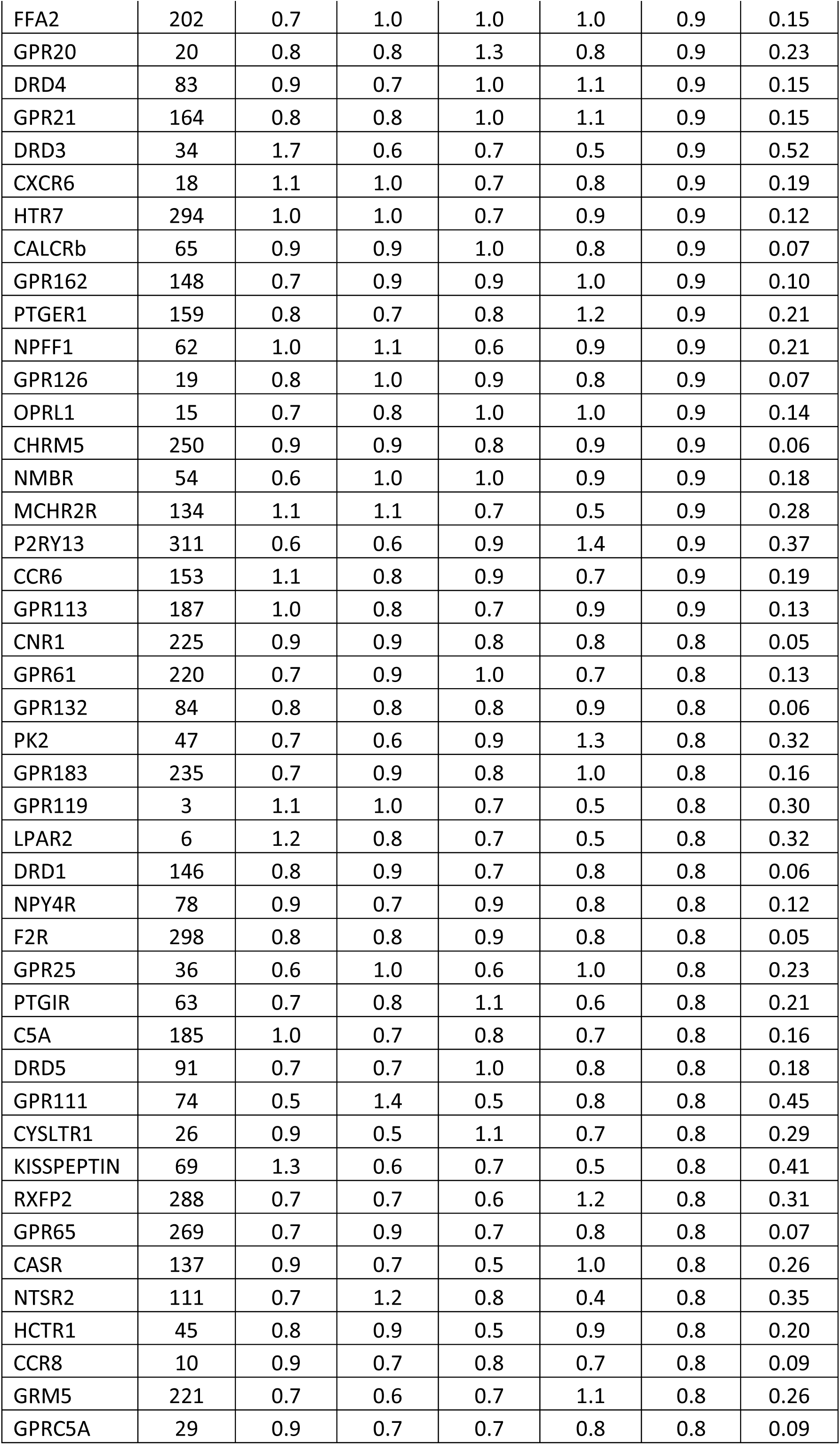
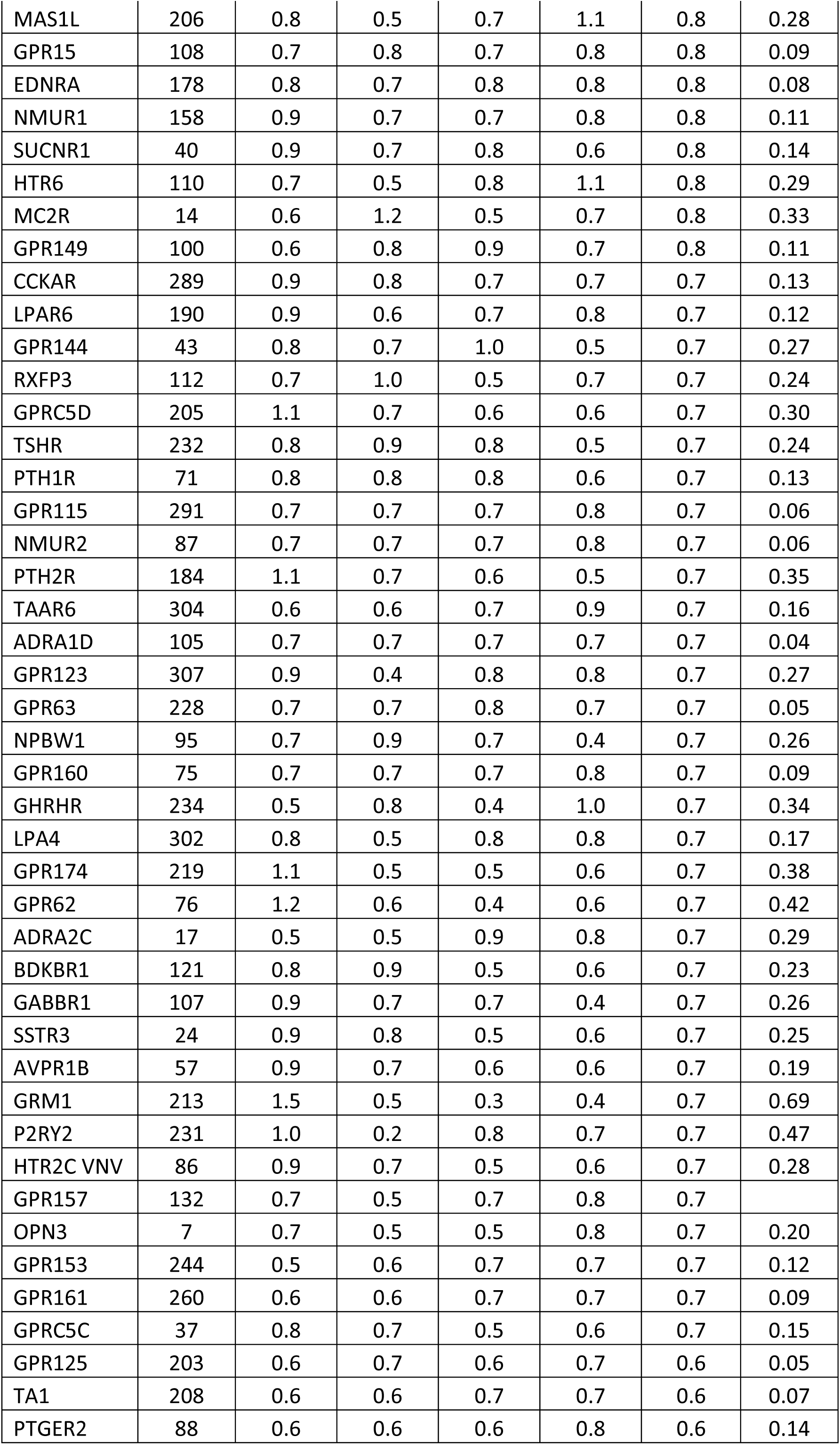
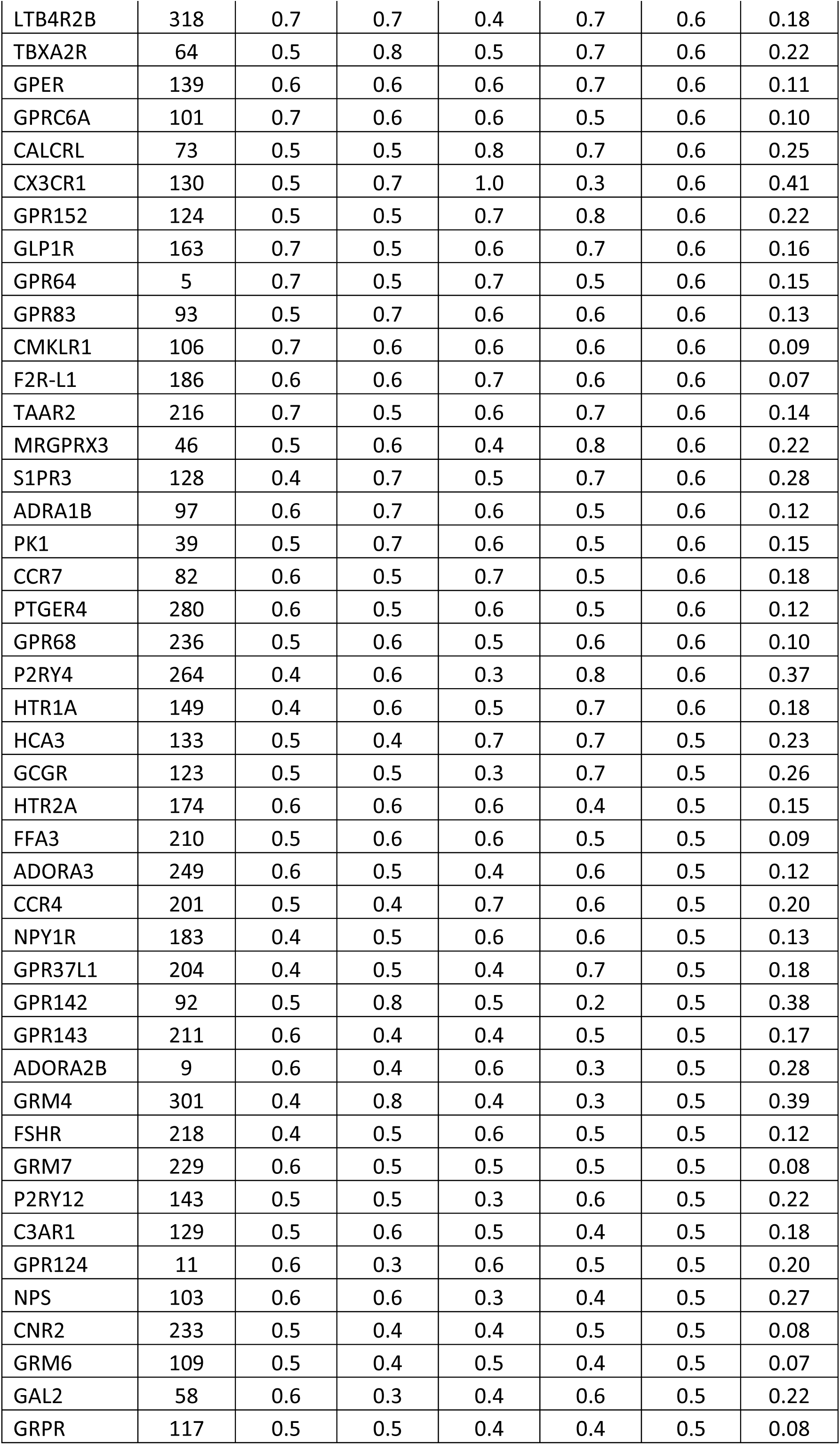
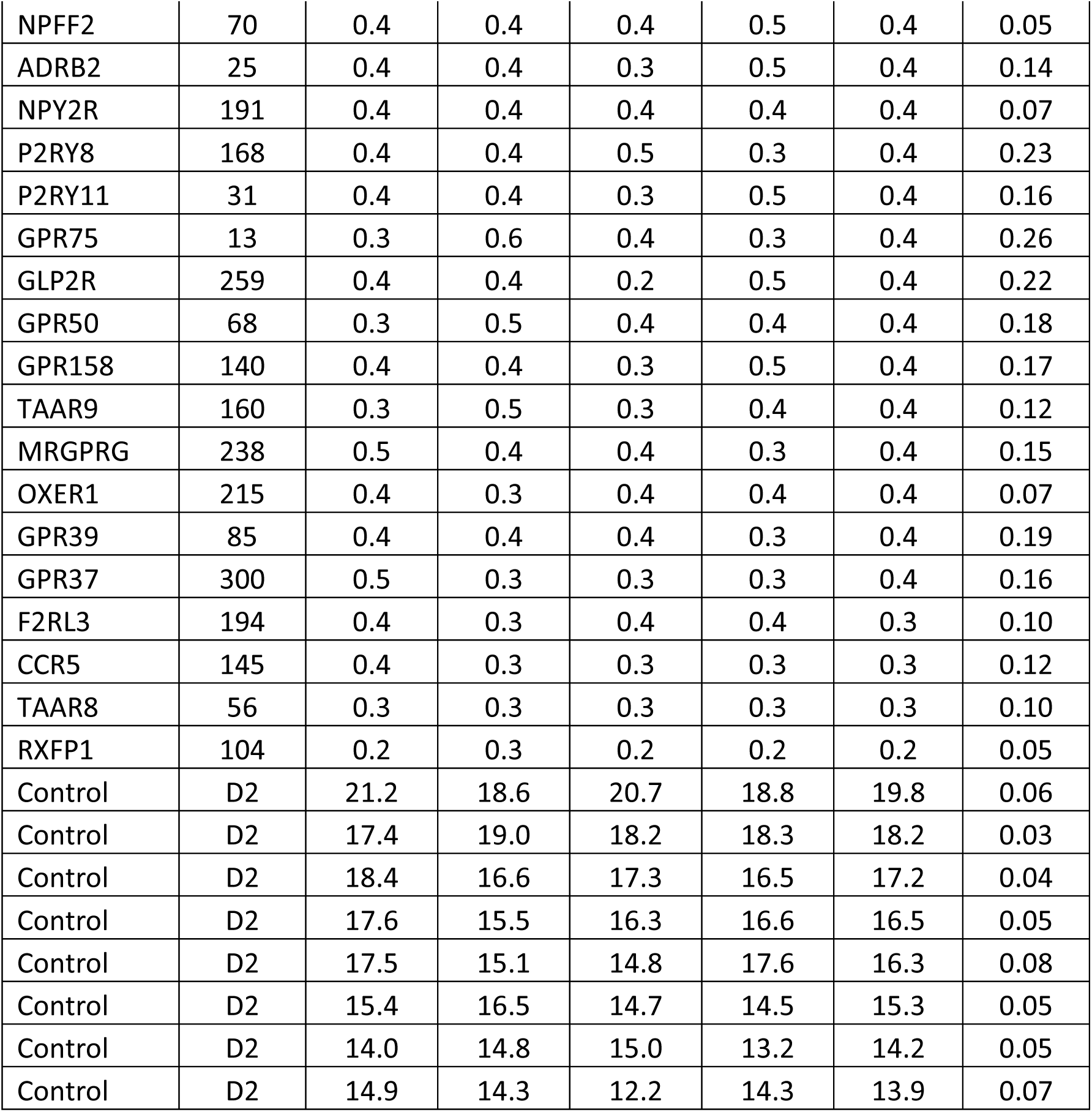
GPCRome screen results with SW016789. Raw fold change data provided by the PDSP core for >300 GPCRs assayed using the Presto-Tango system.

**Table S2.**
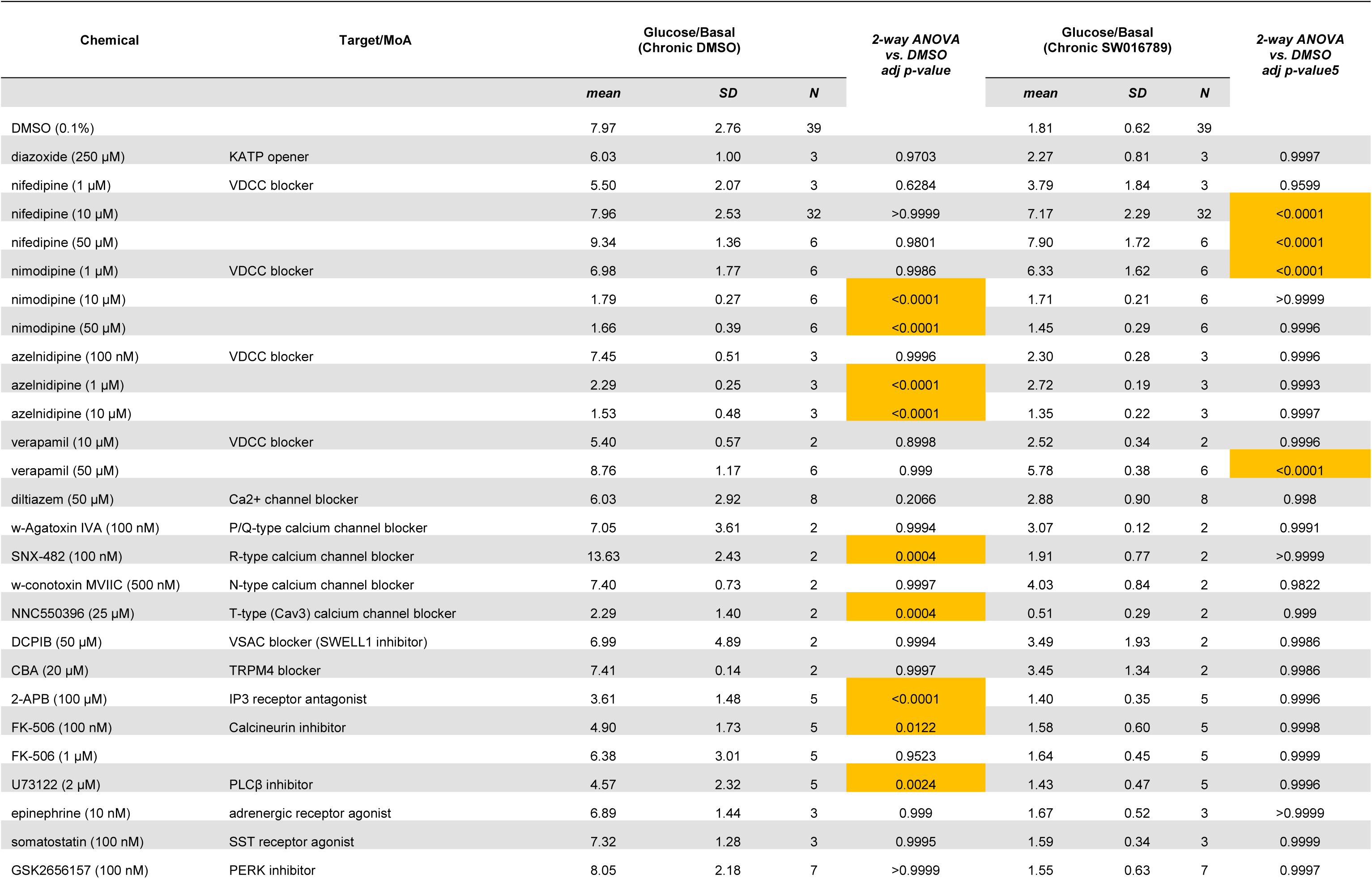
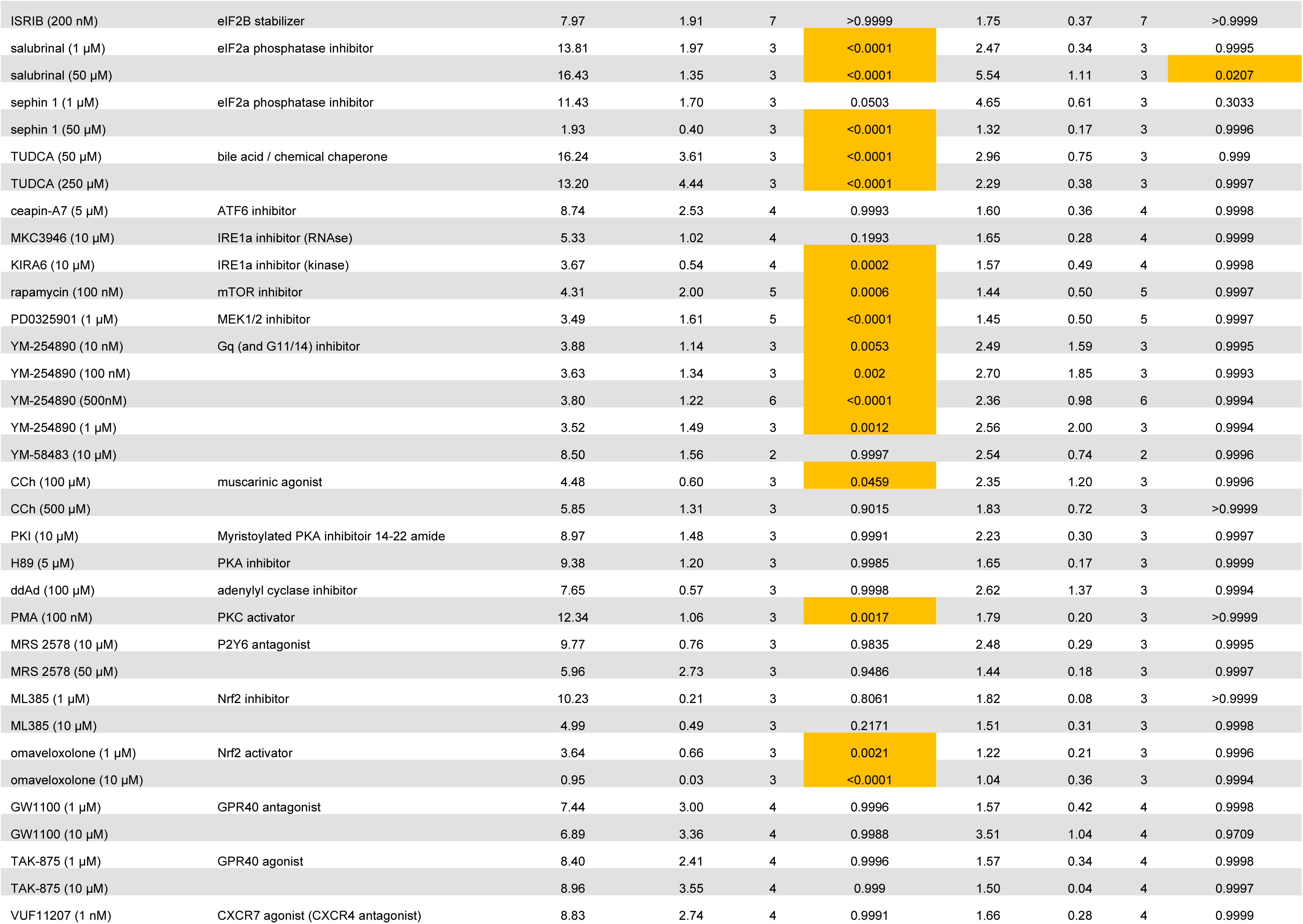
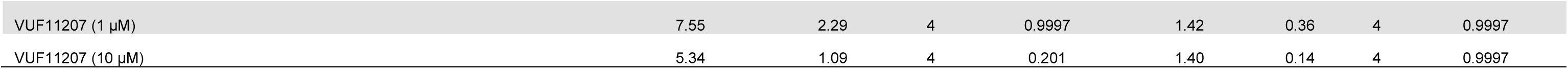
Chemical suppressor screen for mechanism of action studies on SW016789. Fold changes in glucose-stimulated InsGLuc release from MIN6 beta cells pre-exposed to either DMSO or SW016789 (5 µM) for 24 h in the presence or absence of candidate chemical suppressors.

**Table S3.** Raw metabolomics data for SW016789-treated MIN6 cells. Raw total internal current (TIC) values for metabolomics screen.

| HMDB | Pub Chem | KEG G | Metabolite Name | MS Transition | Samples |  |  |  |  |  |  |  |  |  |  |  |
| --- | --- | --- | --- | --- | --- | --- | --- | --- | --- | --- | --- | --- | --- | --- | --- | --- |
|  |  |  |  |  | DMSO_B AS_1 | DMSO_B AS_2 | DMSO_B AS_3 | DMSO_G 30_1 | DMSO_G 30_2 | DMSO_G 30_3 | SW016789_BAS_1 | SW016789_BAS_2 | SW016789_BAS_3 | SW016789_G30_1 | SW016789_G30_2 | SW016789_G30_3 |
| HMDB03331 | 27476 | C02494 | 1-Methyladenosine | 1-methyl-adenosine_pos-1 | 1.29E-03 | 1.16E-03 | 1.04E-03 | 1.20E-03 | 1.51E-03 | 1.55E-03 | 1.54E-03 | 2.14E-03 | 1.72E-03 | 2.13E-03 | 1.69E-03 | 1.54E-03 |
| HMDB00699 | 457 | C02918 | 1-Methylnicotinamide | 1-methyl-nicotinamide_pos-3 | 1.58E-03 | 1.41E-03 | 2.38E-03 | 2.34E-03 | 2.26E-03 | 1.26E-03 | 1.63E-03 | 2.38E-03 | 1.14E-03 | 2.26E-03 | 1.48E-03 | 1.92E-03 |
| HMDB00606 | 439391 | C01087 | 2-Hydroxyglutaric acid | hydroxyglutaric acid_neg-44 | 1.61E-03 | 1.54E-03 | 1.24E-03 | 2.13E-03 | 2.54E-03 | 2.05E-03 | 1.64E-03 | 1.46E-03 | 1.14E-03 | 2.14E-03 | 2.67E-03 | 1.50E-03 |
| HMDB00707 | 979 | C01179 | 4-Hydroxyphenylpyruvic acid | hydroxyphenylpyruvate_pos-73 | 7.77E-04 | 7.28E-04 | 8.60E-04 | 7.92E-04 | 1.16E-03 | 8.42E-04 | 6.71E-04 | 8.08E-04 | 1.04E-03 | 9.90E-04 | 9.31E-04 | 7.84E-04 |
| HMDB00725 | 5810 | C01157 | 4-Hydroxyproline | hydroxy-proline_pos-74 | 3.48E-06 | 3.35E-04 | 4.01E-04 | 4.19E-06 | 3.64E-06 | 6.35E-04 | 3.37E-04 | 3.26E-06 | 3.93E-04 | 3.54E-06 | 3.48E-06 | 4.21E-04 |
| HMDB00017 | 6723 | C00847 | 4-Pyridoxic acid | 4-pyridoxic acid_pos-7 | 1.22E-04 | 1.04E-04 | 2.03E-04 | 2.13E-04 | 1.70E-04 | 8.52E-05 | 1.94E-04 | 2.46E-04 | 2.07E-04 | 1.19E-04 | 1.38E-04 | 4.46E-06 |
| HMDB01173 | 439176 | C00170 | 5'-Methylthioadenosine | S-methyl-5-thioadenosine_pos-134 | 2.69E-02 | 3.47E-02 | 7.43E-02 | 5.00E-02 | 2.66E-02 | 4.75E-02 | 4.49E-02 | 4.72E-02 | 7.18E-02 | 3.89E-02 | 3.67E-02 | 5.76E-02 |
| HMDB01107 | 445404 | - | 7-Methylguanosine | 7-methylguanosine_pos-10 | 3.26E-04 | 5.50E-04 | 3.97E-04 | 3.02E-04 | 2.21E-04 | 2.70E-04 | 1.54E-03 | 1.83E-03 | 1.51E-03 | 1.62E-03 | 2.31E-03 | 1.13E-03 |
| HMDB00895 | 187 | C01996 | Acetylcholine | acetyl-choline_pos-13 | 5.35E-03 | 4.71E-03 | 5.48E-03 | 4.72E-03 | 3.90E-03 | 4.09E-03 | 3.58E-03 | 4.76E-03 | 5.62E-03 | 6.64E-03 | 7.19E-03 | 3.91E-03 |
| HMDB00034 | 190 | C00147 | Adenine | adenine_pos-15 | 1.12E-03 | 8.63E-04 | 8.89E-04 | 8.95E-04 | 1.05E-03 | 8.66E-04 | 1.01E-03 | 1.21E-03 | 1.17E-03 | 1.29E-03 | 1.06E-03 | 1.54E-03 |
| HMDB00050 | 60961 | C00212 | Adenosine | adenosine_pos-16 | 1.53E-02 | 1.56E-02 | 1.87E-02 | 1.51E-02 | 1.97E-02 | 1.49E-02 | 1.15E-02 | 1.96E-02 | 2.40E-02 | 1.96E-02 | 2.60E-02 | 2.79E-02 |
| HMDB00045 | 6083 | C00020 | Adenosine monophosphate | AMP_neg-16 | 3.47E-03 | 3.96E-03 | 3.44E-03 | 1.71E-03 | 1.99E-03 | 1.34E-03 | 2.81E-03 | 3.47E-03 | 4.50E-03 | 1.39E-03 | 1.54E-03 | 1.23E-03 |
| HMDB00019 | 49 | C00141 | Alpha-ketoisovaleric acid | a-ketoisovaleric acid_neg-15 | 3.48E-06 | 3.35E-06 | 5.24E-06 | 1.32E-03 | 1.38E-03 | 1.11E-03 | 3.55E-06 | 3.26E-06 | 5.59E-06 | 1.82E-03 | 1.69E-03 | 1.22E-03 |
| HMDB00510 | 469 | C00956 | Aminoadipic acid | aminoadipic acid_pos-21 | 2.56E-04 | 2.65E-04 | 4.21E-04 | 3.27E-04 | 2.89E-04 | 4.05E-04 | 2.16E-04 | 2.55E-04 | 4.72E-04 | 4.09E-04 | 2.78E-04 | 5.04E-04 |
| HMDB01539 | 123831 | C03626 | Asymmetric dimethylarginine | Ng,Ng-dimethyl-arginine_pos-107 | 1.40E-04 | 3.34E-04 | 2.20E-04 | 6.98E-05 | 2.36E-04 | 9.49E-05 | 2.27E-04 | 2.01E-04 | 1.21E-04 | 1.82E-04 | 9.33E-05 | 1.07E-04 |
| HMDB00043 | 247 | C00719 | Betaine | betaine_pos-26 | 1.79E-02 | 1.69E-02 | 2.69E-02 | 2.55E-02 | 2.62E-02 | 4.59E-02 | 2.11E-02 | 1.89E-02 | 3.09E-02 | 2.55E-02 | 2.66E-02 | 2.90E-02 |
| HMDB02013 | 439829 | C02862 | Butyrylcarnitine | carnitine-C4_pos-37 | 1.33E-03 | 1.19E-03 | 2.47E-04 | 1.66E-03 | 3.18E-03 | 4.16E-04 | 1.09E-03 | 1.18E-03 | 2.22E-04 | 2.73E-03 | 2.08E-03 | 4.73E-04 |
| HMDB0011671 | NA | NA | cADP-ribose | cADP-ribose_neg-18 | 1.59E-03 | 1.60E-03 | 1.60E-03 | 1.85E-03 | 1.79E-03 | 2.32E-03 | 1.66E-03 | 1.12E-03 | 1.96E-03 | 1.76E-03 | 1.37E-03 | 1.47E-03 |
| HMDB01096 | 278 | C00169 | Carbamoyl phosphate | carbamoylphosphate_neg-19 | 2.10E-04 | 2.64E-04 | 1.28E-04 | 2.52E-04 | 2.03E-04 | 9.13E-05 | 2.53E-04 | 2.85E-04 | 1.26E-04 | 2.51E-04 | 2.40E-04 | 5.62E-05 |
| HMDB00062 | 10917 | C00487 | Carnitine | carnitine_pos-30 | 1.81E-02 | 2.15E-02 | 3.00E-02 | 3.15E-02 | 2.77E-02 | 2.80E-02 | 2.03E-02 | 1.67E-02 | 1.90E-02 | 3.05E-02 | 2.55E-02 | 1.94E-02 |
| HMDB00097 | 305 | C00114 | Choline | choline_pos-41 | 1.43E-03 | 2.01E-03 | 1.26E-03 | 1.24E-03 | 1.43E-03 | 8.61E-04 | 1.91E-03 | 2.23E-03 | 2.97E-03 | 1.11E-03 | 1.64E-03 | 1.55E-03 |
| HMDB00094 | 311 | C00158 | Citric acid | citrate-isocitrate_neg-22 | 3.60E-02 | 4.22E-02 | 3.85E-02 | 5.24E-02 | 6.90E-02 | 6.60E-02 | 3.47E-02 | 3.64E-02 | 2.88E-02 | 4.82E-02 | 5.80E-02 | 6.00E-02 |
| HMDB00064 | 586 | C00300 | Creatine | creatine_pos-44 | 6.25E-03 | 9.22E-03 | 1.02E-02 | 4.52E-03 | 5.71E-03 | 5.63E-03 | 6.56E-03 | 6.38E-03 | 1.02E-02 | 4.13E-03 | 3.91E-03 | 4.18E-03 |
| HMDB00562 | 588 | C00791 | Creatinine | creatinine_pos-45 | 1.09E-04 | 3.21E-04 | 2.99E-04 | 9.49E-04 | 5.86E-04 | 7.41E-04 | 1.17E-04 | 1.58E-04 | 1.81E-04 | 4.70E-04 | 2.96E-04 | 5.02E-04 |
| HMDB00089 | 6175 | C00475 | Cytidine | cytidine_pos-49 | 9.49E-04 | 1.28E-03 | 1.98E-03 | 1.06E-03 | 6.18E-04 | 1.16E-03 | 1.60E-03 | 3.37E-03 | 1.04E-03 | 1.87E-03 | 1.64E-03 | 2.02E-03 |
| HMDB00095 | 6131 | C00055 | Cytidine monophosphate | CMP_pos-43 | 1.92E-03 | 1.08E-03 | 9.29E-04 | 1.39E-03 | 1.61E-03 | 1.36E-03 | 1.26E-03 | 1.34E-03 | 6.96E-04 | 1.25E-03 | 9.23E-04 | 8.16E-04 |
| HMDB00905 | 12599 | C00360 | Deoxyadenosine monophosphate | dAMP_pos-51 | 7.20E-04 | 1.24E-03 | 3.24E-03 | 1.04E-03 | 8.90E-04 | 2.41E-03 | 7.83E-04 | 9.28E-04 | 1.69E-03 | 2.10E-03 | 3.28E-03 | 1.51E-04 |
| HMDB00014 | 137<br>11 | C008<br>81 | Deoxycytidine | 2-deoxycytidine_pos-6 | 5.95E-05 | 7.59E-05 | 1.62E-04 | 1.14E-04 | 1.16E-04 | 1.48E-04 | 8.54E-05 | 1.27E-04 | 5.43E-05 | 9.66E-05 | 1.05E-04 | 1.20E-04 |
| HMDB00071 | 650<br>58 | C055<br>12 | Deoxyinosine | deoxyinosine_pos-52 | 3.48E-06 | 9.76E-05 | 7.02E-05 | 6.10E-05 | 1.38E-04 | 5.26E-06 | 1.17E-04 | 1.29E-04 | 1.94E-04 | 2.67E-04 | 1.42E-04 | 1.96E-04 |
| HMDB01351 | 439<br>287 | C006<br>72 | Deoxyribose 1-phosphate | deoxyribose-phosphate_neg-23 | 5.66E-04 | 5.12E-04 | 9.71E-04 | 6.06E-04 | 4.48E-04 | 9.17E-04 | 4.14E-04 | 5.29E-04 | 1.10E-03 | 3.72E-04 | 6.18E-04 | 8.30E-04 |
| HMDB00122 | 579<br>3 | C000<br>31 | D-Glucose | glucose-fructose_neg-36 | 3.48E-06 | 2.79E-05 | 5.24E-06 | 4.22E-04 | 4.28E-04 | 2.60E-04 | 3.28E-05 | 2.71E-05 | 5.59E-06 | 3.72E-04 | 3.83E-04 | 3.59E-04 |
| HMDB00127 | 444<br>791 | C001<br>91 | D-Glucuronic acid | glucuronate_neg-38 | 3.23E-04 | 2.68E-04 | 1.83E-04 | 4.38E-04 | 5.71E-04 | 5.13E-04 | 2.93E-04 | 2.95E-04 | 1.67E-04 | 4.55E-04 | 4.47E-04 | 5.34E-04 |
| HMDB03339 | 233<br>27 | C002<br>17 | D-Glutamic acid | glutamate_neg-39 | 1.68E-02 | 1.58E-02 | 1.77E-02 | 1.73E-02 | 1.20E-02 | 1.60E-02 | 1.73E-02 | 1.50E-02 | 1.58E-02 | 1.25E-02 | 1.14E-02 | 1.34E-02 |
| HMDB01473 | 668 | C001<br>11 | Dihydroxyacetone phosphate | dihydroxyacetone-phosphate_neg-25 | 2.40E-04 | 3.19E-04 | 6.49E-04 | 4.57E-04 | 3.54E-04 | 7.18E-04 | 3.25E-04 | 2.58E-04 | 6.51E-04 | 4.04E-04 | 3.39E-04 | 5.25E-04 |
| HMDB00092 | 673 | C010<br>26 | Dimethylglycine | dimethylglycine_pos-54 | 2.85E-03 | 2.73E-03 | 2.44E-03 | 1.93E-03 | 1.97E-03 | 1.65E-03 | 2.97E-03 | 2.42E-03 | 3.24E-03 | 2.28E-03 | 2.22E-03 | 5.63E-03 |
| HMDB01548 | 439<br>167 | C001<br>17 | D-Ribose 5-phosphate | ribose 5-phosphate_neg-66 | 1.11E-04 | 1.18E-04 | 2.25E-04 | 4.19E-06 | 3.64E-06 | 1.46E-04 | 1.14E-04 | 8.75E-05 | 2.40E-04 | 8.19E-05 | 3.48E-06 | 4.46E-06 |
| HMDB01068 | 228<br>335<br>59 | C002<br>81 | D-Sedoheptulose 7-phosphate | sedoheptulose-7-phosphate_neg-68 | 1.99E-04 | 2.80E-04 | 2.60E-04 | 2.50E-04 | 1.71E-04 | 2.91E-04 | 2.17E-04 | 2.11E-04 | 3.28E-04 | 1.84E-04 | 2.63E-04 | 2.85E-04 |
| HMDB00121 | 603<br>7 | C005<br>04 | Folic acid | folic acid_pos-57 | 3.76E-04 | 3.94E-04 | 2.98E-04 | 3.04E-04 | 4.12E-04 | 2.38E-04 | 4.42E-04 | 3.68E-04 | 4.26E-04 | 2.96E-04 | 3.22E-04 | 2.85E-04 |
| HMDB01058 | 172<br>313 | C003<br>54 | Fructose 1,6-bisphosphate | fructose-1,6-bisphosphate_neg-27 | 3.48E-06 | 2.17E-05 | 4.12E-05 | 2.67E-04 | 3.33E-04 | 4.57E-04 | 3.55E-06 | 3.26E-06 | 6.72E-05 | 3.37E-04 | 3.80E-04 | 2.17E-04 |
| HMDB00645 | 123<br>912 | C004<br>46 | Galactose 1-phosphate | galactose 1-phosphate_neg-30 | 1.24E-04 | 1.22E-04 | 6.78E-05 | 1.11E-04 | 1.75E-04 | 2.17E-04 | 1.18E-04 | 9.65E-05 | 1.68E-04 | 1.32E-04 | 1.40E-04 | 4.33E-05 |
| HMDB00112 | 119 | C003<br>34 | Gamma-Aminobutyric acid | γ-aminobutyric acid_pos-155 | 5.12E-03 | 6.72E-03 | 5.94E-03 | 4.67E-03 | 6.56E-03 | 5.73E-03 | 5.61E-03 | 7.00E-03 | 1.16E-02 | 6.85E-03 | 7.08E-03 | 9.32E-03 |
| HMDB00625 | 106<br>90 | C002<br>57 | Gluconic acid | gluconic acid_neg-34 | 1.24E-04 | 4.72E-05 | 1.67E-04 | 6.97E-05 | 3.64E-06 | 1.14E-04 | 9.85E-05 | 3.26E-06 | 1.60E-04 | 3.76E-05 | 3.48E-06 | 4.46E-06 |
| HMDB01401 | 595<br>8 | C000<br>92 | Glucose 6-phosphate | G6P-F6P_neg-29 | 6.75E-05 | 5.11E-05 | 7.02E-05 | 5.70E-04 | 3.53E-04 | 6.55E-04 | 2.79E-05 | 6.33E-05 | 3.62E-05 | 4.13E-04 | 3.88E-04 | 4.09E-04 |
| HMDB00125 | 124<br>886 | C000<br>51 | Glutathione | GSH_pos-64 | 3.12E-02 | 1.26E-02 | 2.02E-02 | 1.64E-02 | 1.51E-02 | 3.01E-02 | 1.67E-02 | 1.31E-02 | 2.71E-02 | 1.64E-02 | 1.69E-02 | 1.88E-02 |
| HMDB00126 | 439<br>162 | C000<br>93 | Glycerol 3-phosphate | sn-glycerol-3-phosphate_neg-69 | 2.11E-04 | 2.23E-04 | 4.02E-04 | 1.90E-04 | 1.09E-04 | 1.48E-04 | 3.51E-04 | 2.35E-04 | 4.60E-04 | 1.47E-04 | 1.67E-04 | 1.39E-04 |
| HMDB00086 | 719<br>20 | C006<br>70 | Glycerophosphocholine | glycerophosphorylcholine_pos-61 | 8.38E-02 | 7.55E-02 | 4.72E-02 | 5.81E-02 | 5.82E-02 | 4.07E-02 | 6.16E-02 | 6.49E-02 | 2.65E-02 | 6.85E-02 | 6.22E-02 | 3.76E-02 |
| HMDB00123 | 750 | C000<br>37 | Glycine | glycine_pos-62 | 1.82E-04 | 2.06E-04 | 2.94E-04 | 1.57E-04 | 1.55E-04 | 1.29E-04 | 1.58E-04 | 1.67E-04 | 1.47E-04 | 1.32E-04 | 1.01E-04 | 1.90E-04 |
| HMDB00119 | 760 | C000<br>48 | Glyoxylic acid | glyoxylate_neg-42 | 3.48E-06 | 3.35E-06 | 5.24E-06 | 2.46E-04 | 2.66E-04 | 1.80E-04 | 3.28E-05 | 3.26E-06 | 7.75E-05 | 3.34E-04 | 1.51E-04 | 3.47E-04 |
| HMDB00128 | 763 | C005<br>81 | Guanidoacetic acid | guanidoacetic acid_pos-66 | 1.98E-03 | 2.26E-03 | 1.10E-03 | 2.53E-03 | 2.66E-03 | 1.01E-03 | 2.48E-03 | 1.68E-03 | 1.53E-03 | 1.89E-03 | 2.37E-03 | 1.54E-03 |
| HMDB00132 | 764 | C002<br>42 | Guanine | guanine_pos-67 | 9.78E-04 | 1.16E-03 | 6.25E-04 | 6.40E-04 | 6.79E-04 | 2.26E-04 | 1.08E-03 | 9.94E-04 | 5.19E-04 | 6.16E-04 | 6.08E-04 | 2.29E-04 |
| HMDB00133 | 680<br>2 | C003<br>87 | Guanosine | guanosine_pos-68 | 5.85E-03 | 9.17E-03 | 3.39E-03 | 1.06E-02 | 5.31E-03 | 6.52E-03 | 1.60E-02 | 1.69E-02 | 6.75E-03 | 2.11E-02 | 2.03E-02 | 1.45E-02 |
| HMDB00157 | 790 | C002<br>62 | Hypoxanthine | hypoxanthine_pos-75 | 2.26E-03 | 1.08E-03 | 2.47E-03 | 1.13E-03 | 1.43E-03 | 8.83E-04 | 2.06E-03 | 1.76E-03 | 3.27E-03 | 1.62E-03 | 2.14E-03 | 1.04E-03 |
| HMDB00738 | 798 | C004<br>63 | Indole | indole_pos-79 | 7.86E-04 | 8.57E-04 | 4.96E-04 | 8.10E-04 | 8.64E-04 | 4.82E-04 | 7.39E-04 | 7.31E-04 | 5.21E-04 | 7.99E-04 | 9.18E-04 | 5.93E-04 |
| HMDB00195 | 602<br>1 | C002<br>94 | Inosine | inosine_pos-90 | 3.89E-03 | 2.62E-03 | 1.43E-03 | 4.59E-03 | 3.53E-03 | 9.84E-03 | 1.17E-02 | 8.08E-03 | 3.66E-03 | 7.77E-03 | 7.33E-03 | 7.64E-03 |
| HMDB00175 | 858<br>2 | C001<br>30 | Inosinic acid | IMP_pos-78 | 1.19E-02 | 1.59E-02 | 1.43E-02 | 5.27E-03 | 6.57E-03 | 5.42E-03 | 1.03E-02 | 1.12E-02 | 9.58E-03 | 3.80E-03 | 4.45E-03 | 4.65E-03 |
| HMDB00688 | 642<br>685<br>1 | - | Isovalerylcarnitine | carnitine-C5_pos-38 | 7.73E-04 | 1.77E-03 | 3.56E-04 | 1.60E-03 | 2.93E-03 | 7.98E-04 | 1.09E-03 | 1.33E-03 | 4.24E-04 | 3.16E-03 | 2.08E-03 | 7.10E-04 |
| HMDB00748 | 643<br>327 | C056<br>07 | L-3-Phenyllactic acid | phenyl-lactic acid_neg-58 | 1.04E-03 | 8.35E-04 | 1.50E-03 | 1.11E-03 | 1.31E-03 | 1.10E-03 | 1.22E-03 | 8.36E-04 | 2.02E-03 | 9.50E-04 | 1.23E-03 | 1.26E-03 |
| HMDB00201 | 1 | C025<br>71 | L-Acetylcarnitine | acetyl-carnitine_pos-12 | 1.71E-01 | 1.24E-01 | 1.52E-01 | 1.31E-01 | 9.27E-02 | 1.39E-01 | 1.83E-01 | 1.60E-01 | 1.38E-01 | 1.35E-01 | 1.05E-01 | 1.57E-01 |
| HMDB00161 | 595<br>0 | C000<br>41 | L-Alanine | alanine_pos-19 | 7.81E-04 | 7.40E-04 | 1.38E-03 | 9.96E-04 | 8.70E-04 | 1.39E-03 | 7.05E-04 | 6.51E-04 | 1.25E-03 | 7.72E-04 | 7.99E-04 | 1.04E-03 |
| HMDB00517 | 632<br>2 | C000<br>62 | L-Arginine | arginine_pos-22 | 3.46E-03 | 3.39E-03 | 2.56E-03 | 2.16E-03 | 2.21E-03 | 1.27E-03 | 2.08E-03 | 2.09E-03 | 1.89E-03 | 1.81E-03 | 1.50E-03 | 8.87E-04 |
| HMDB00168 | 626<br>7 | C001<br>52 | L-Asparagine | asparagine_pos-23 | 1.01E-02 | 9.13E-03 | 5.46E-03 | 1.07E-02 | 9.93E-03 | 6.17E-03 | 8.87E-03 | 8.14E-03 | 6.31E-03 | 8.93E-03 | 1.02E-02 | 8.41E-03 |
| HMDB00191 | 596<br>0 | C000<br>49 | L-Aspartic acid | aspartate_pos-24 | 2.09E-03 | 1.79E-03 | 2.84E-03 | 1.12E-03 | 9.93E-04 | 1.64E-03 | 1.17E-03 | 1.65E-03 | 2.65E-03 | 1.00E-03 | 6.72E-04 | 1.35E-03 |
| HMDB00099 | 439<br>258 | C022<br>91 | L-Cystathionine | cystathionine_pos-46 | 1.24E-03 | 8.77E-04 | 1.51E-03 | 2.26E-03 | 1.56E-03 | 1.81E-03 | 1.75E-03 | 1.49E-03 | 1.28E-03 | 1.43E-03 | 1.15E-03 | 1.59E-03 |
| HMDB00574 | 586<br>2 | C000<br>97 | L-Cysteine | cysteine_pos-47 | 6.43E-05 | 9.14E-05 | 8.72E-05 | 4.19E-06 | 3.64E-06 | 5.26E-06 | 6.40E-05 | 1.01E-04 | 8.53E-05 | 4.26E-05 | 3.48E-06 | 4.46E-06 |
| HMDB00641 | 596<br>1 | C000<br>64 | L-Glutamine | glutamine_pos-60 | 1.42E-03 | 1.95E-03 | 7.78E-04 | 1.39E-03 | 2.82E-03 | 1.23E-03 | 1.11E-03 | 1.19E-03 | 9.24E-04 | 1.20E-03 | 2.39E-03 | 9.97E-04 |
| HMDB00756 | 324<br>693<br>8 | - | L-Hexanoylcarnitine | carnitine-C6_pos-39 | 3.40E-03 | 2.70E-03 | 2.70E-03 | 1.83E-03 | 1.37E-03 | 1.21E-03 | 3.94E-03 | 2.77E-03 | 3.38E-03 | 2.33E-03 | 1.98E-03 | 9.37E-04 |
| HMDB00177 | 627<br>4 | C001<br>35 | L-Histidine | histidine_pos-69 | 1.44E-04 | 2.06E-04 | 5.24E-06 | 1.47E-04 | 1.63E-04 | 1.36E-04 | 1.24E-04 | 1.53E-04 | 1.49E-04 | 1.72E-04 | 1.77E-04 | 1.28E-04 |
| HMDB00719 | 126<br>47 | C002<br>63 | L-Homoserine | homoserine_pos-72 | 9.73E-04 | 1.09E-03 | 7.48E-04 | 7.71E-04 | 8.08E-04 | 6.78E-04 | 9.36E-04 | 8.29E-04 | 5.12E-04 | 8.11E-04 | 7.80E-04 | 7.93E-04 |
| HMDB00172 | 630<br>6 | C004<br>07 | L-Isoleucine | isoleucine_pos-91 | 8.79E-02 | 1.09E-01 | 5.70E-02 | 9.63E-02 | 1.04E-01 | 5.65E-02 | 9.13E-02 | 9.36E-02 | 6.73E-02 | 1.06E-01 | 9.98E-02 | 7.69E-02 |
| HMDB00190 | 107<br>689 | C001<br>86 | L-Lactic acid | lactate_neg-48 | 3.08E-04 | 2.78E-04 | 7.05E-04 | 2.69E-03 | 2.52E-03 | 3.09E-03 | 2.24E-04 | 1.67E-04 | 8.35E-04 | 3.09E-03 | 2.41E-03 | 4.02E-03 |
| HMDB00182 | 596<br>2 | C000<br>47 | L-Lysine | lysine_pos-96 | 3.45E-03 | 3.98E-03 | 2.43E-03 | 1.42E-03 | 1.36E-03 | 1.04E-03 | 2.22E-03 | 2.05E-03 | 1.48E-03 | 1.14E-03 | 1.88E-03 | 9.41E-04 |
| HMDB00156 | 222<br>656 | C001<br>49 | L-Malic acid | malic acid_neg-49 | 3.48E-06 | 3.35E-06 | 5.24E-06 | 7.70E-04 | 8.63E-04 | 6.01E-04 | 3.55E-06 | 3.26E-06 | 5.59E-06 | 8.76E-04 | 8.73E-04 | 6.54E-04 |
| HMDB00696 | 613<br>7 | C000<br>73 | L-Methionine | methionine_pos-98 | 2.45E-03 | 3.34E-03 | 1.60E-03 | 3.39E-03 | 4.07E-03 | 2.23E-03 | 3.61E-03 | 3.40E-03 | 1.62E-03 | 3.27E-03 | 3.35E-03 | 2.28E-03 |
| HMDB00791 | 119<br>538<br>14 | C028<br>38 | L-Octanoylcarnitine | carnitine-C8_pos-40 | 1.98E-03 | 2.37E-03 | 1.11E-03 | 2.29E-03 | 1.62E-03 | 6.47E-04 | 2.32E-03 | 2.16E-03 | 1.26E-03 | 1.41E-03 | 1.23E-03 | 1.09E-03 |
| HMDB00159 | 614<br>0 | C000<br>79 | L-Phenylalanine | phenyl-alanine_pos-113 | 1.66E-02 | 1.87E-02 | 1.06E-02 | 2.51E-02 | 2.29E-02 | 9.32E-03 | 1.91E-02 | 1.44E-02 | 1.20E-02 | 2.44E-02 | 1.82E-02 | 1.49E-02 |
| HMDB00162 | 145<br>742 | C001<br>48 | L-Proline | proline_pos-118 | 6.03E-02 | 7.42E-02 | 5.60E-02 | 5.31E-02 | 8.30E-02 | 5.27E-02 | 5.26E-02 | 6.46E-02 | 4.56E-02 | 5.04E-02 | 5.87E-02 | 3.92E-02 |
| HMDB00187 | 595<br>1 | C000<br>65 | L-Serine | serine_pos-130 | 6.94E-04 | 5.16E-04 | 7.63E-04 | 1.25E-03 | 6.35E-04 | 6.45E-04 | 4.95E-04 | 7.68E-04 | 1.12E-03 | 6.07E-04 | 8.32E-04 | 5.80E-04 |
| HMDB00158 | 605<br>7 | C000<br>82 | L-Tyrosine | tyrosine_pos-146 | 8.80E-03 | 8.59E-03 | 8.35E-03 | 1.17E-02 | 1.28E-02 | 6.85E-03 | 1.00E-02 | 1.13E-02 | 1.05E-02 | 8.35E-03 | 1.32E-02 | 1.05E-02 |
| HMDB00883 | 628<br>7 | C001<br>83 | L-Valine | valine_pos-152 | 1.84E-03 | 1.90E-03 | 1.36E-03 | 1.74E-03 | 2.43E-03 | 1.28E-03 | 1.73E-03 | 1.99E-03 | 1.04E-03 | 1.95E-03 | 1.79E-03 | 1.51E-03 |
| HMDB02005 | 847 | - | Methionine sulfoxide | methionine sulfoxide_pos-97 | 2.96E-04 | 3.45E-04 | 3.85E-04 | 3.98E-04 | 4.11E-04 | 3.99E-04 | 3.14E-04 | 4.69E-04 | 3.49E-04 | 3.36E-04 | 3.35E-04 | 3.08E-04 |
| HMDB00211 | 892 | C001<br>37 | Myo-inositol | myo-inositol_pos-99 | 2.12E-04 | 4.25E-04 | 4.53E-04 | 3.68E-04 | 4.63E-04 | 2.90E-04 | 3.38E-04 | 3.76E-04 | 3.41E-04 | 3.45E-04 | 3.55E-04 | 6.11E-04 |
| HMDB01325 | 440<br>120 | C037<br>93 | N6,N6,N6-Trimethyl-L-lysine | trimethyllysine_pos-144 | 2.87E-03 | 3.44E-03 | 2.15E-03 | 3.70E-03 | 3.83E-03 | 1.68E-03 | 2.17E-03 | 2.51E-03 | 1.68E-03 | 2.74E-03 | 2.36E-03 | 1.47E-03 |
| HMDB00215 | 439<br>174 | C001<br>40 | N-Acetyl-D-glucosamine | N-acetyl-D-glucosamine 1-phosphate_neg-54 | 1.42E-04 | 1.37E-04 | 2.03E-04 | 7.69E-05 | 8.10E-05 | 8.89E-05 | 2.55E-04 | 1.10E-04 | 2.44E-04 | 3.95E-05 | 6.86E-05 | 6.39E-05 |
| HMDB00766 | 880<br>64 | C006<br>24 | N-Acetyl-L-alanine | N-acetyl-alanine_neg-53 | 5.41E-04 | 3.77E-04 | 5.84E-04 | 6.59E-04 | 5.15E-04 | 9.46E-04 | 2.69E-04 | 3.74E-04 | 4.53E-04 | 7.20E-04 | 3.34E-04 | 3.43E-04 |
| HMDB00812 | 650<br>65 | C010<br>42 | N-Acetyl-L-aspartic acid | N-acetyl-aspartic acid_pos-101 | 7.25E-04 | 5.25E-04 | 2.82E-04 | 2.88E-04 | 6.03E-04 | 8.21E-04 | 5.29E-04 | 7.96E-04 | 1.11E-03 | 7.02E-04 | 6.50E-04 | 4.66E-04 |
| HMDB00902 | 589<br>3 | C000<br>03 | NAD | NAD_pos-105 | 1.30E-01 | 1.43E-01 | 1.65E-01 | 1.57E-01 | 1.52E-01 | 1.83E-01 | 1.24E-01 | 1.27E-01 | 1.70E-01 | 1.33E-01 | 1.61E-01 | 1.49E-01 |
| HMDB00446 | 192<br>590 | C129<br>89 | N-Alpha-acetyllysine | acetyllysine_pos-14 | 2.68E-03 | 2.57E-03 | 1.74E-03 | 2.13E-03 | 2.81E-03 | 1.48E-03 | 2.05E-03 | 2.06E-03 | 1.54E-03 | 1.63E-03 | 2.18E-03 | 1.05E-03 |
| HMDB01406 | 936 | C001<br>53 | Niacinamide | nicotinamide_pos-108 | 7.08E-04 | 1.05E-03 | 7.10E-04 | 1.35E-03 | 1.58E-03 | 2.21E-03 | 9.36E-04 | 1.00E-03 | 3.02E-04 | 1.25E-03 | 8.50E-04 | 7.52E-04 |
| HMDB00214 | 626<br>2 | C000<br>77 | Ornithine | ornithine_pos-111 | 2.95E-04 | 2.53E-04 | 3.54E-04 | 1.82E-04 | 2.15E-04 | 1.37E-04 | 1.44E-04 | 1.98E-04 | 1.99E-04 | 1.39E-04 | 9.97E-05 | 1.35E-04 |
| HMDB03337 | 975 | C001<br>27 | Oxidized glutathione | GSSG_pos-65 | 1.30E-03 | 3.36E-03 | 1.99E-03 | 6.08E-04 | 1.37E-03 | 1.34E-03 | 1.32E-03 | 1.76E-03 | 3.00E-03 | 9.54E-04 | 5.73E-04 | 1.37E-03 |
| HMDB00208 | 51 | C000<br>26 | Oxoglutaric acid | a-keto-glutarate_neg-14 | 1.60E-04 | 2.11E-04 | 5.24E-06 | 5.20E-04 | 8.14E-04 | 5.54E-04 | 9.69E-05 | 1.50E-04 | 5.59E-06 | 5.41E-04 | 6.65E-04 | 3.54E-04 |
| HMDB00210 | 988 | C00864 | Pantothenic acid | pantothenic_acid_pos-112 | 2.79E-02 | 2.46E-02 | 2.39E-02 | 3.43E-02 | 3.08E-02 | 2.47E-02 | 2.21E-02 | 2.54E-02 | 2.00E-02 | 2.90E-02 | 2.55E-02 | 3.64E-02 |
| HMDB01511 | 587 | C02305 | Phosphocreatine | phospho-creatine_neg-60 | 1.11E-04 | 1.24E-04 | 1.59E-04 | 4.31E-04 | 3.87E-04 | 4.40E-04 | 8.54E-05 | 1.55E-04 | 8.01E-05 | 3.00E-04 | 3.04E-04 | 3.81E-04 |
| HMDB01565 | 1014 | C00588 | Phosphorylcholine | phosphorylcholine_pos-115 | 2.33E-02 | 2.01E-02 | 4.01E-02 | 1.73E-02 | 2.16E-02 | 3.22E-02 | 2.31E-02 | 1.73E-02 | 4.40E-02 | 2.38E-02 | 1.27E-02 | 4.70E-02 |
| HMDB00070 | 849 | C00408 | Pipecolic acid | pipecolic_acid_pos-116 | 7.90E-03 | 5.44E-03 | 5.08E-03 | 6.95E-03 | 8.67E-03 | 6.26E-03 | 6.46E-03 | 6.33E-03 | 4.31E-03 | 6.98E-03 | 5.91E-03 | 5.32E-03 |
| HMDB00824 | 107738 | C03017 | Propionylcarnitine | carnitine-C3_pos-36 | 3.31E-02 | 2.93E-02 | 9.61E-03 | 9.61E-03 | 1.31E-02 | 3.39E-03 | 3.09E-02 | 4.00E-02 | 2.48E-02 | 1.62E-02 | 2.16E-02 | 4.66E-03 |
| HMDB13305 | 75612 | - | Propionylcholine | propionylcholine_pos-119 | 1.08E-02 | 1.14E-02 | 1.13E-02 | 1.27E-02 | 1.31E-02 | 1.32E-02 | 1.09E-02 | 1.20E-02 | 1.11E-02 | 1.10E-02 | 1.27E-02 | 1.04E-02 |
| HMDB01414 | 1045 | C00134 | Putrescine | putrescine_pos-121 | 1.32E-04 | 1.73E-04 | 2.42E-04 | 1.69E-04 | 1.82E-04 | 1.70E-04 | 1.77E-04 | 1.69E-04 | 2.51E-04 | 1.08E-04 | 4.04E-04 | 1.03E-04 |
| HMDB01431 | 1052 | C00534 | Pyridoxamine | pyridoxamine_pos-122 | 1.11E-04 | 1.78E-04 | 5.24E-06 | 2.65E-04 | 3.58E-04 | 2.94E-04 | 9.19E-05 | 1.06E-04 | 5.59E-06 | 2.37E-04 | 3.62E-04 | 1.97E-04 |
| HMDB00232 | 1066 | C03722 | Quinolinic acid_pos-125 | quinolinate_pos-125 | 4.40E-04 | 2.35E-04 | 6.94E-04 | 9.00E-04 | 7.32E-04 | 1.15E-03 | 4.10E-04 | 3.69E-04 | 4.70E-04 | 6.97E-04 | 5.15E-04 | 6.65E-04 |
| HMDB00244 | 493570 | C00255 | Riboflavin | riboflavin_pos-127 | 2.85E-04 | 2.43E-04 | 3.08E-04 | 3.49E-04 | 3.62E-04 | 2.07E-04 | 8.84E-04 | 6.92E-04 | 5.45E-04 | 4.77E-04 | 8.98E-04 | 3.18E-04 |
| HMDB00939 | 439155 | C00021 | S-Adenosylhomocysteine | S-adenosyl-homocysteine_neg-67 | 6.63E-04 | 7.63E-04 | 1.64E-03 | 6.92E-04 | 5.99E-04 | 1.04E-03 | 1.09E-03 | 9.23E-04 | 2.70E-03 | 5.65E-04 | 5.60E-04 | 1.11E-03 |
| HMDB01185 | 16757548 | C00019 | S-Adenosylmethionine | S-adenosyl-methionine_pos-128 | 7.40E-03 | 1.21E-02 | 1.07E-02 | 8.25E-03 | 1.86E-02 | 1.81E-02 | 1.22E-02 | 8.92E-03 | 1.51E-02 | 8.63E-03 | 9.65E-03 | 1.86E-02 |
| HMDB03070 | 8742 | C00493 | Shikimic acid | shikimic_acid_pos-131 | 1.13E-02 | 8.70E-03 | 9.38E-03 | 6.23E-03 | 6.24E-03 | 4.53E-03 | 6.70E-03 | 8.18E-03 | 5.65E-03 | 5.03E-03 | 4.05E-03 | 3.11E-03 |
| HMDB01066 | 440018 | C03451 | S-Lactoylglutathione | S-lactoylglutathione_pos-133 | 3.48E-06 | 3.35E-06 | 5.24E-06 | 1.84E-03 | 1.83E-03 | 3.00E-03 | 3.55E-06 | 3.26E-06 | 5.59E-06 | 2.01E-03 | 1.18E-03 | 2.20E-03 |
| HMDB01257 | 1102 | C00315 | Spermidine | spermidine_pos-135 | 8.12E-03 | 1.34E-02 | 1.22E-02 | 8.27E-03 | 1.24E-02 | 1.42E-02 | 1.23E-02 | 1.40E-02 | 2.50E-02 | 1.51E-02 | 1.75E-02 | 1.44E-02 |
| HMDB01256 | 1103 | C00750 | Spermine | spermine_pos-136 | 1.44E-03 | 1.78E-03 | 3.24E-03 | 5.26E-03 | 3.15E-03 | 3.81E-03 | 3.29E-03 | 4.43E-03 | 2.81E-03 | 2.25E-03 | 7.24E-03 | 1.71E-03 |
| HMDB00848 | 6426855 | - | Stearoylcarnitine | carnitine-C18_pos-34 | 9.05E-04 | 7.81E-04 | 4.19E-04 | 3.64E-04 | 1.89E-04 | 1.73E-04 | 9.93E-04 | 9.43E-04 | 4.96E-04 | 3.90E-04 | 1.22E-04 | 2.02E-04 |
| HMDB00254 | 1110 | C00042 | Succinic acid | succinate_neg-70 | 2.54E-03 | 2.71E-03 | 2.16E-03 | 2.27E-03 | 3.48E-03 | 3.62E-03 | 2.82E-03 | 2.38E-03 | 2.26E-03 | 2.73E-03 | 4.66E-03 | 2.69E-03 |
| HMDB00251 | 1123 | C00245 | Taurine | taurine_neg-71 | 2.67E-02 | 2.29E-02 | 3.47E-02 | 2.65E-02 | 2.33E-02 | 2.79E-02 | 2.73E-02 | 2.17E-02 | 3.17E-02 | 2.34E-02 | 2.11E-02 | 2.52E-02 |
| HMDB05066 | 53477791 | - | Tetradecanoylcarnitine | carnitine-C14_pos-32 | 7.75E-04 | 1.47E-03 | 5.06E-04 | 2.77E-04 | 3.20E-04 | 2.80E-04 | 6.72E-04 | 1.11E-03 | 4.65E-04 | 4.04E-04 | 1.33E-04 | 3.08E-04 |
| HMDB00235 | 1130 | C00378 | Thiamine | thiamine_pos-139 | 5.75E-04 | 6.64E-04 | 4.65E-04 | 5.81E-04 | 5.41E-04 | 3.83E-04 | 3.66E-04 | 5.01E-04 | 3.67E-04 | 5.55E-04 | 6.79E-04 | 2.29E-04 |
| HMDB00273 | 5789 | C00214 | Thymidine | thymidine_pos-142 | 6.75E-05 | 9.91E-05 | 5.24E-06 | 4.19E-06 | 7.24E-05 | 5.26E-06 | 7.64E-05 | 1.12E-04 | 6.46E-05 | 1.26E-04 | 1.20E-04 | 4.46E-06 |
| HMDB00300 | 1174 | C00106 | Uracil | uracil_pos-147 | 1.83E-04 | 9.91E-05 | 8.72E-05 | 6.20E-05 | 3.64E-06 | 9.00E-05 | 1.13E-04 | 9.65E-05 | 1.65E-04 | 3.54E-06 | 6.92E-05 | 6.40E-05 |
| HMDB00296 | 6029 | C00299 | Uridine | uridine_neg-77 | 3.48E-06 | 3.25E-05 | 5.24E-06 | 2.71E-05 | 2.19E-05 | 5.11E-05 | 6.40E-05 | 3.77E-05 | 2.07E-05 | 3.77E-05 | 4.02E-05 | 4.33E-05 |
| HMDB00295 | 6031 | C00015 | Uridine 5'-diphosphate | UDP_neg-72 | 8.68E-05 | 8.98E-05 | 7.26E-05 | 1.09E-04 | 9.93E-05 | 8.52E-05 | 8.37E-05 | 5.73E-05 | 1.45E-04 | 1.24E-04 | 3.59E-04 | 7.43E-05 |
| HMDB00292 | 1188 | C00385 | Xanthine | xanthine_neg-78 | 2.80E-04 | 3.55E-04 | 1.69E-04 | 2.50E-04 | 2.71E-04 | 1.07E-04 | 3.25E-04 | 3.20E-04 | 2.92E-04 | 2.42E-04 | 2.61E-04 | 1.44E-04 |
| HMDB00299 | 64959 | C01762 | Xanthosine | xanthosine_pos-154 | 4.16E-04 | 4.56E-04 | 1.79E-04 | 6.49E-04 | 5.54E-04 | 7.28E-04 | 3.28E-04 | 4.41E-04 | 3.62E-05 | 4.42E-04 | 1.07E-03 | 5.18E-04 |

**Table S4.**
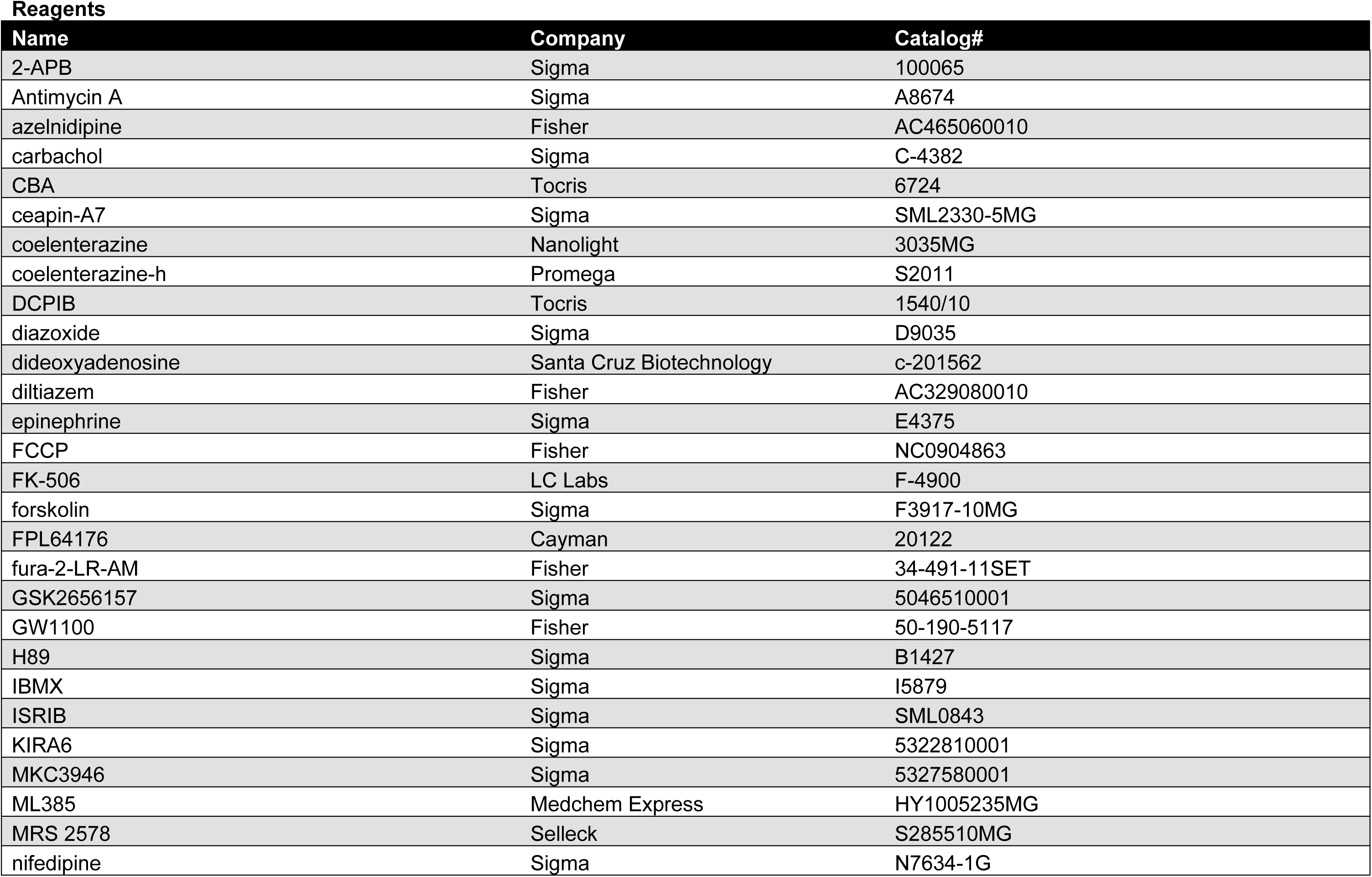
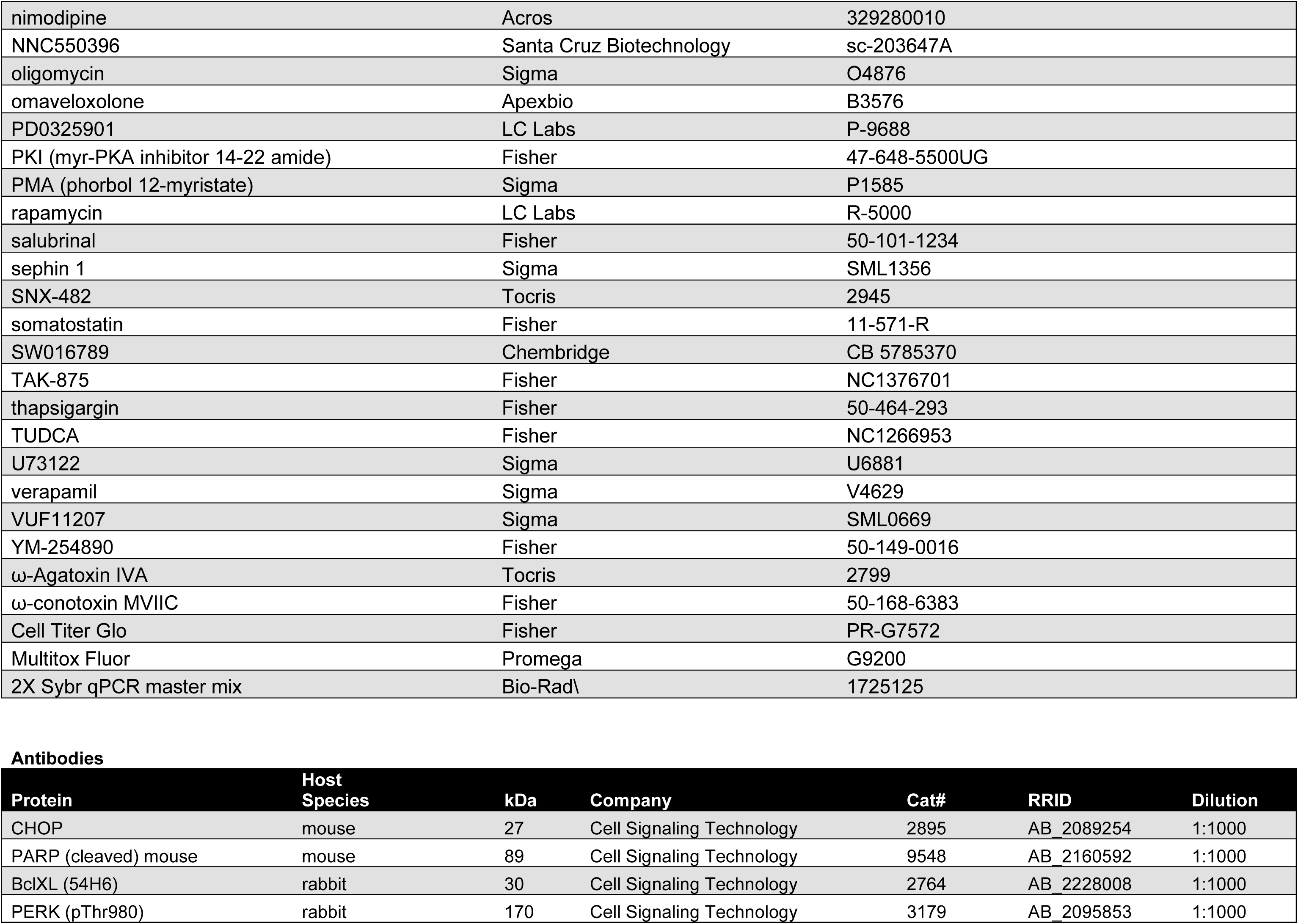
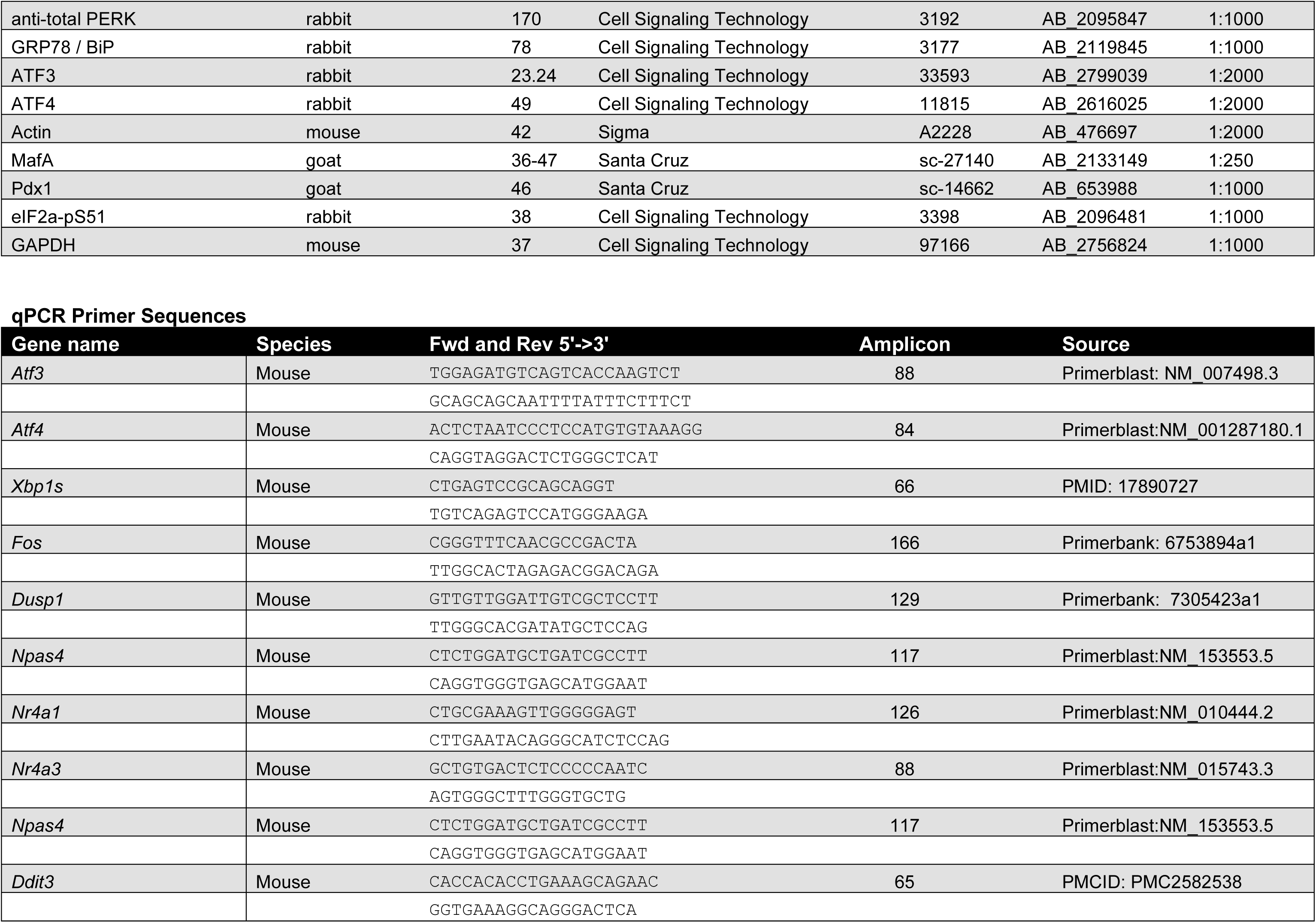
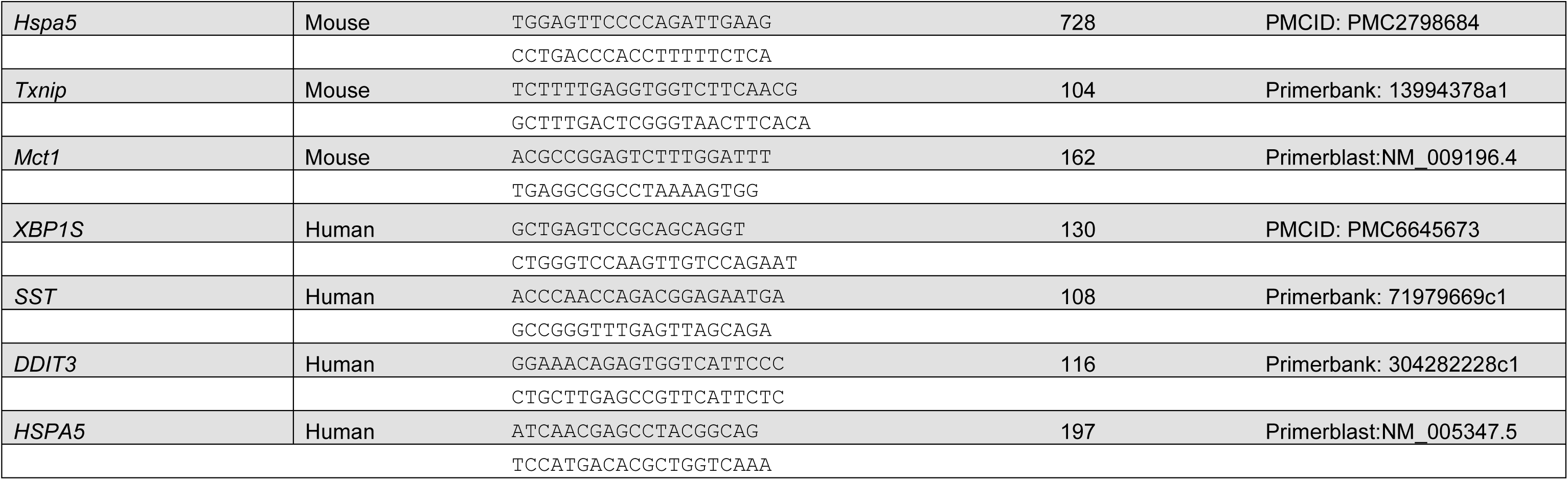
Research material sources. All information pertaining to chemical reagents, antibodies, qPCR primers, and donor human islets.

## Checklist for reporting human islet preparations used in research

Adapted from Hart NJ, Powers AC (2018) Progress, challenges, and suggestions for using human islets to understand islet biology and human diabetes. Diabetologia https://doi.org/10.1007/s00125-018-4772-2

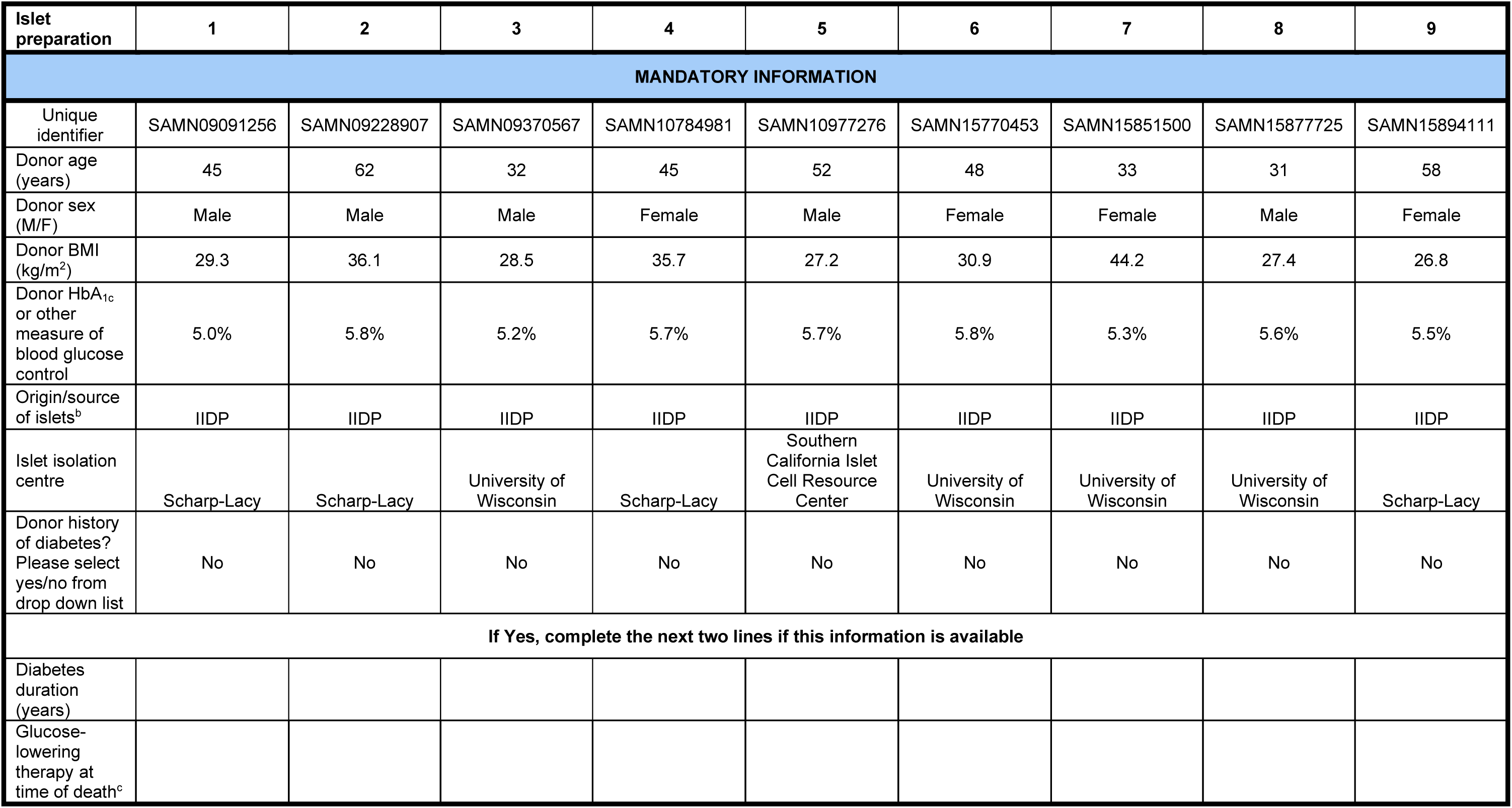
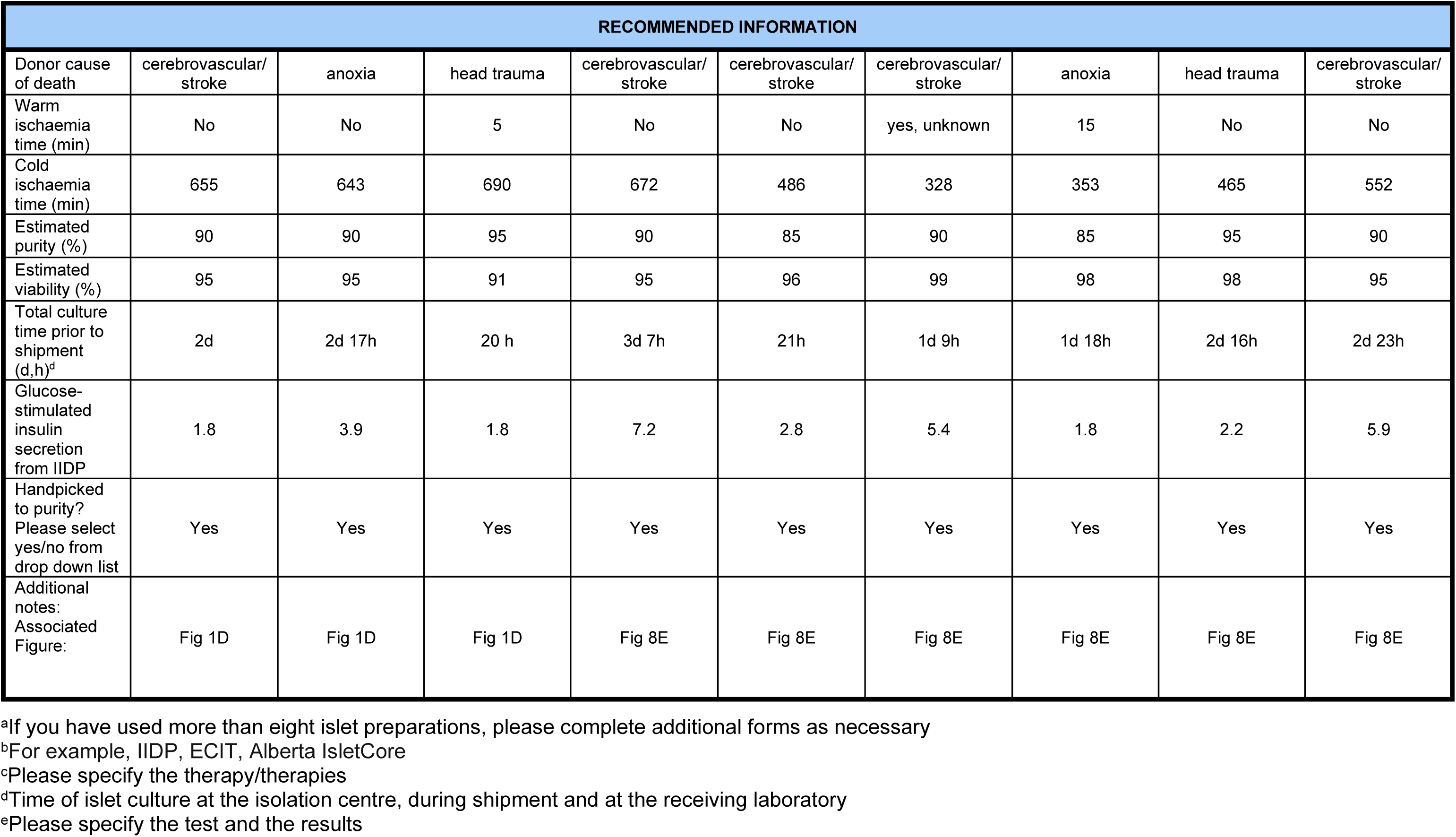

